# A suicidal and extensively disordered luciferase with a bright luminescence

**DOI:** 10.1101/2023.12.21.572513

**Authors:** Fenne Marjolein Dijkema, Marta Iglesia Escarpizo-Lorenzana, Matilde Knapkøien Nordentoft, Hanna Christin Rabe, Cagla Sahin, Michael Landreh, Rui Mamede Branca, Esben Skipper Sørensen, Brian Søndergaard Christensen, Andreas Prestel, Kaare Teilum, Jakob Rahr Winther

**Affiliations:** The Linderstrøm-Lang Centre for Protein Science; Department of Biology; University of Copenhagen; Denmark; Department of Microbiology, Tumor and Cell Biology; Karolinska Institutet; Stockholm; Sweden; Science for Life Laboratory; Department of Oncology-Pathology; Karolinska Institutet; Stockholm; Sweden; Department of Molecular Biology and Genetics; Section for Cellular Health, Intervention and Nutrition; Aarhus University; Denmark

## Abstract

Gaussia luciferase (GLuc) is one of the most luminescent luciferases known and is widely used as a reporter in biochemistry and cell biology. During catalysis GLuc undergoes inactivation by irreversible covalent modification. The mechanism by which GLuc generates luminescence and how it becomes inactivated are however not known. Here we show that GLuc unlike other enzymes has an extensively disordered structure with a minimal hydrophobic core and no apparent binding pocket for the main substrate, coelenterazine. From an alanine scan, we identified two Arg residues required for light production. These residues separated with an average of about 22 Å and a major structural rearrangement is required if they are to interact with the substrate simultaneously. We furthermore show that in addition to coelenterazine, GLuc also can oxidize furimazine, however, in this case without production of light. Both substrates result in the formation of adducts with the enzyme, which eventually leads to enzyme inactivation. Our results demonstrate that a rigid protein structure and substrate binding site are no prerequisites for high enzymatic activity and specificity. In addition to the increased understanding of enzymes in general, the findings will facilitate future improvement of GLuc as a reporter luciferase.

**Significance statement:** Enzymes are typically characterized by an overall globular structure with a hydrophobic core and a defined cavity for binding of substrate, containing the active site amino acid residues. Gaussia Luciferase is a widely used luminescent reporter with a very strong, albeit short-lived, flash of light due to rapid auto-inactivation. We show, using solution NMR, that while this luciferase shows some secondary structure elements held together by disulfide bonds this highly unusual enzyme is extensively disordered with essentially no hydrophobic core. Although the enzymatic mechanism remains unknown, we have identified two essential arginine residues but, in the structure, these do not point into a common active site. In spite of this, the enzyme has high substrate specificity suggesting that it undergoes major structural rearrangements upon binding of substrate.

## Introduction

Most enzymes are globular proteins with well-defined structures, densely packed hydrophobic cores and substrate binding pockets with catalytic residues at precise positions. Intrinsically disordered enzymes, on the other hand, are exceedingly rare. There are only few examples of enzymes with extensive structural dynamics. The protein UreG, for instance, a GTPase responsible for delivering Ni^2+^ to nickel-dependent urease, is disordered in its apo state but folds upon binding the urease accessory proteins UreF and UreD to form a well-structured and active complex (Palombo et al., 2017). Perhaps the best example of an enzyme that preserves activity while being intrinsically disordered, is an engineered version of the enzyme chorismate mutase from *Methanococcus jannaschii* (Vamvaca et al., 2004). Here the native enzyme is a stable well-structured homodimer where a long α-helix spans the dimer. In an attempt to allow the helix to bend back and generate a monomer, a flexible linker was inserted in this helix. Surprisingly, this construct had properties of a molten globule while preserving essentially full enzymatic activity. Thus, enzymes can be fully active catalysts while being highly dynamic. However, most examples of highly dynamic or molten globule enzymes are engineered in one way or another (Uversky, 2023).

Gaussia luciferase (GLuc) is the luciferase from the copepod *Gaussia princeps*. It catalyzes the oxidative decarboxylation of coelenterazine (CTZ) into coelenteramide (CTM). Excited coelenteramide then emits a photon at 480 nm (Verhaegen & Christopoulos, 2002). This light-emitting reaction is extremely bright, with a sensitivity down to femtomolar concentration of the luciferase (Larionova et al., 2018). In its native host, the protein is secreted (Bowlby & Case, 1991) and thus the gene includes an N-terminal signal peptide of 18 residues. After cleavage of this signal peptide, the protein contains 168 amino acid residues and has a molecular mass of 18.2 kDa. Because of this small size, GLuc has become a popular bioluminescence-based reporter in life sciences. Due to the presence of 5 essential disulfide bridges, it is mostly used as a reporter in the secretory pathway or extracellularly. GLuc has likely arisen from a gene duplication of a domain of approximately 70 amino acids (Inouye & Sahara, 2008), which has resulted in a conserved spacing of four cysteine residues in each domain (Supplementary Figure S1).

Because of its use as a reporter, many studies have focused on improving bioluminescence characteristics and developed enhanced GLuc variants. With two goals in particular: 1) to shift the emission wavelength from 480 nm to a longer wavelength that is less absorbed by biological tissues (Kim et al., 2011; Wu et al., 2016) and 2) to prolong the light signal duration (Kim et al., 2011; Welsh et al., 2009). We have recently investigated GLuc’s luminescence and shown that it fades exponentially with a substrate concentration-dependent half-life ranging from 15 seconds to a few minutes. We showed that the decay was caused by an autocatalytic inactivation after less than 200 reaction cycles. Inactivation was accompanied by an increase in size of the protein, apparent by SDS-PAGE and size exclusion chromatography. It also caused the appearance of a fluorescence signal at 410 nm (Dijkema et al, 2021).

A structure of GLuc has recently been proposed based on nuclear magnetic resonance (NMR) spectroscopy (PDB ID: 7D2O; Wu et al., 2020). The protein used was a variant featuring two mutations, E100A and G103R, which were introduced to increase the expression yield. Large regions of the structure were only defined by few and local NOE restraints. Even the most structured parts of the structure were not well-packed, with large surface-accessible areas. The main part of the structure consists of four helices in a double V-shape. The most N-terminal helix was positioned inside the V-cleft, perpendicular to the four main helices. Three disulfide bridges held the structure together at the kink of the V. Two further disulfide bridges stabilized two small helices at either open end of the V. A CTZ binding site was proposed between one side of the V cleft and the N-terminal helix.

Prior to the publication of the aforementioned structure, GLuc served as a structure prediction target in the 2020 Critical Assessment of protein Structure Prediction (CASP14) (Kryshtafovych et al., 2021). It proved a very difficult target to predict. The best-scoring model in the competition, AlphaFold2 (Jumper et al., 2021), achieved a Global distance test (GDT) score of 61.11 (out of max. 100), while its mean GDT score across all targets was 92.4. Only 6 out of 110 prediction targets received lower scores than GLuc, suggesting unusual properties of this protein. Indeed, the NMR-derived structure 7D2O does not resemble any previously published structures (Wu et al., 2020).

The mechanism of GLuc is thus far unresolved. Even though CTZ has been known as a luciferin for decades, it was only recently that a mechanism of CTZ-dependent luciferases was proposed. This mechanism was based on the structure of Renilla luciferase in complex with a substrate analogue (Schenkmayerova et al., 2023). Nevertheless, Renilla luciferase and GLuc share no sequence relationship and thus GLuc does not share this discovered mechanism.

To shed light on GLuc’s reaction mechanism and its substrate-dependent inactivation, we identified active site residues in GLuc by an alanine scan of conserved polar amino acid residues. Since the results of this scan were not compatible with the published structure (which was based on a mutant variant), we revisited the structural analysis of GLuc. While the overall structure had similar helices in a double V-shape, we saw no sign of a central helix in the V-cleft. We also found a different disulfide pattern through mass spectrometry. Nevertheless, this new structure is also not compatible with location of essential amino acid residues identified by the alanine scan. However, it did suggest a high level of dynamics, which was supported by low protection factors in time-resolved Hydrogen Deuterium eXchange (HDX) as determined by NMR.

Our results show that GLuc is a highly disordered enzyme that despite a minimal hydrophobic core and no apparent substrate binding site retains a high catalytic activity.

## Results and discussion

### Alanine scan reveals two amino acid residues essential for function

To map the active site of GLuc, we first set out to identify critical catalytic residues, which were not obvious from the 7D2O structure. We performed an alanine scan based on two assumptions: 1) important residues would be conserved among homologous enzymes, and 2) such residues should be able to form hydrogen bonds to be relevant as catalytic entities and promote chemical reactions per se. To identify conserved residues, we used the POSSUM server (Wang et al., 2017) to determine residue conservation scores in GLuc compared to similar sequences from the UniRef50 database (Steinegger & Söding, 2018; Suzek et al., 2015).

POSSUM considers in the conservation score that some residues can functionally replace each other by using a position specific scoring matrix (PSSM). The core helixes received slightly higher conservation scores than the more unstructured parts, especially around the bottom of the V-shaped structure. Interestingly, the N-terminal helix received relatively low conservation scores throughout from POSSUM. To gauge the relevance of the 24 N-terminal residues we prepared and purified variants in which these were deleted. Truncation of the first 23 amino acids resulted in a drop of activity to about 3.5 % of the activity of the full-length protein. The activity decay rate also increased about 2-fold (SI Figure S5). The C-terminus of GLuc, contained several partly conserved residues, namely glycine 164 and 167 and aspartate 168, even though this part of the structure appears disordered and no long-range distance restraints were found (Wu et al., 2020). To address this apparent disparity, we prepared enzyme variants truncated at the C-terminal by 4, 13 and 17 residues, resulting in variants with 19 %, 1.4 % and 0.13 % of the activity of full length GLuc, respectively (Figure 1c and SI Figure S3). The activity of GLuc thus decreases gradually with progressive truncations, supporting the notion that both N- and C-termini play functional roles without being essential. Notably, all deletions yielded stable and soluble proteins.

**Figure 1.**
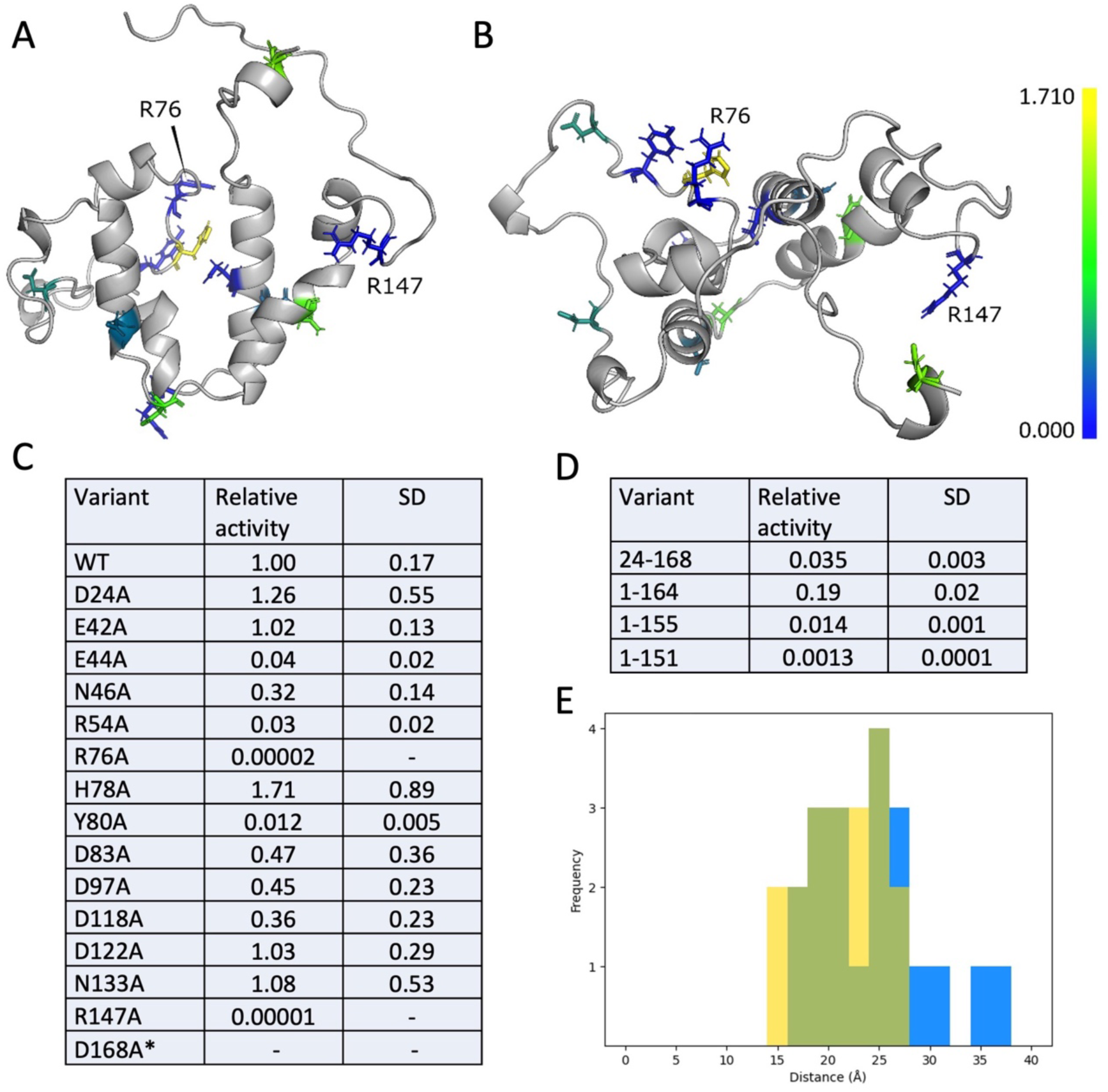
Structure and alanine scan of GLuc. Solution structure determined by NMR (PDB: 8OQB), seen from the front (**A**) and top (**B**) with conserved residues included in the alanine scan colored from least (blue) and most conserved (yellow) calculated using POSSUM (Wang et al., 2017). **C**) Alanine substitution and N- and C-terminal truncation variants of GLuc and their activities relative to wild-type GLuc. R27A did not express well enough for characterization and although E42A and D168A did display activity the expression yield was too low for accurate determination of specific activity. **D**) Activity of truncation mutants relative to wild type. **E**) Comparison of the distance between residues R76 and R147 in the structure 7D2O (Wu et al. (2020)) (yellow) and 8OQB (blue) with overlap shown in green. In both, the distribution of distances shows that the residues are too far apart to simultaneously contact CTZ.

Using a cutoff at a POSSUM conservation score of 0.7, we selected 16 residues with the ability to form hydrogen bonds for an Ala-scan (excluding cysteine, since all cysteines in GLuc are involved in disulfide bonds). Because of the rapid decay in activity of GLuc during catalytic turnover, substrate was injected directly into the stirred cuvette and initial rates were assessed as luminescence output. Initial rates at *t* = 0 were estimated by extrapolation of the intensity curve, using an in-house developed Python package (https://github.com/FDijkema/LumParser). In addition, we determined decay rate constants for all variants (Supplementary Table S2).

Most variants were fairly active with only four variants having rates at less than 10% of the reference. Of these R76 and R147 stood out as being essentially inactive. There are several possible roles for arginine in the active site. 1) An arginine has been shown to coordinate oxygen in Renilla luciferase (Schenkmayerova et al., 2023). However, as we will show later, CTZ is still oxidized and reacts with the protein, in the absence of these arginines in GLuc. 2) The arginines stabilize the cation intermediate in the oxidation of CTZ, as hypothesized for an artificially designed luciferase recently reported (Yeh et al., 2023). Finally, 3) the arginines participate in substrate binding. Since CTZ contains a large aromatic system, a positively charged side chain could favorably interact with the π-electron cloud of one of its rings, either through a π-cation interaction or via a stacking with the planar guanidino-group (Kumar et al., 2018).

CTZ produces luminescence through reaction with oxygen on its own when dissolved in organic solvent (Lucas & Solano, 1992) and one could argue that GLuc merely provides a hydrophobic environment for CTZ to bind in a non-specific way, as suggested by the hydrophobic patch seen in the V cleft in Figure 2C. However, GLuc has a very high substrate-specificity for CTZ, while Furimazine (FRZ) or quite a few other luciferin analogues (Inouye et al., 2013) are very poor substrates. To investigate this apparent paradox, we revisited the GLuc structure.

**Figure 2.**
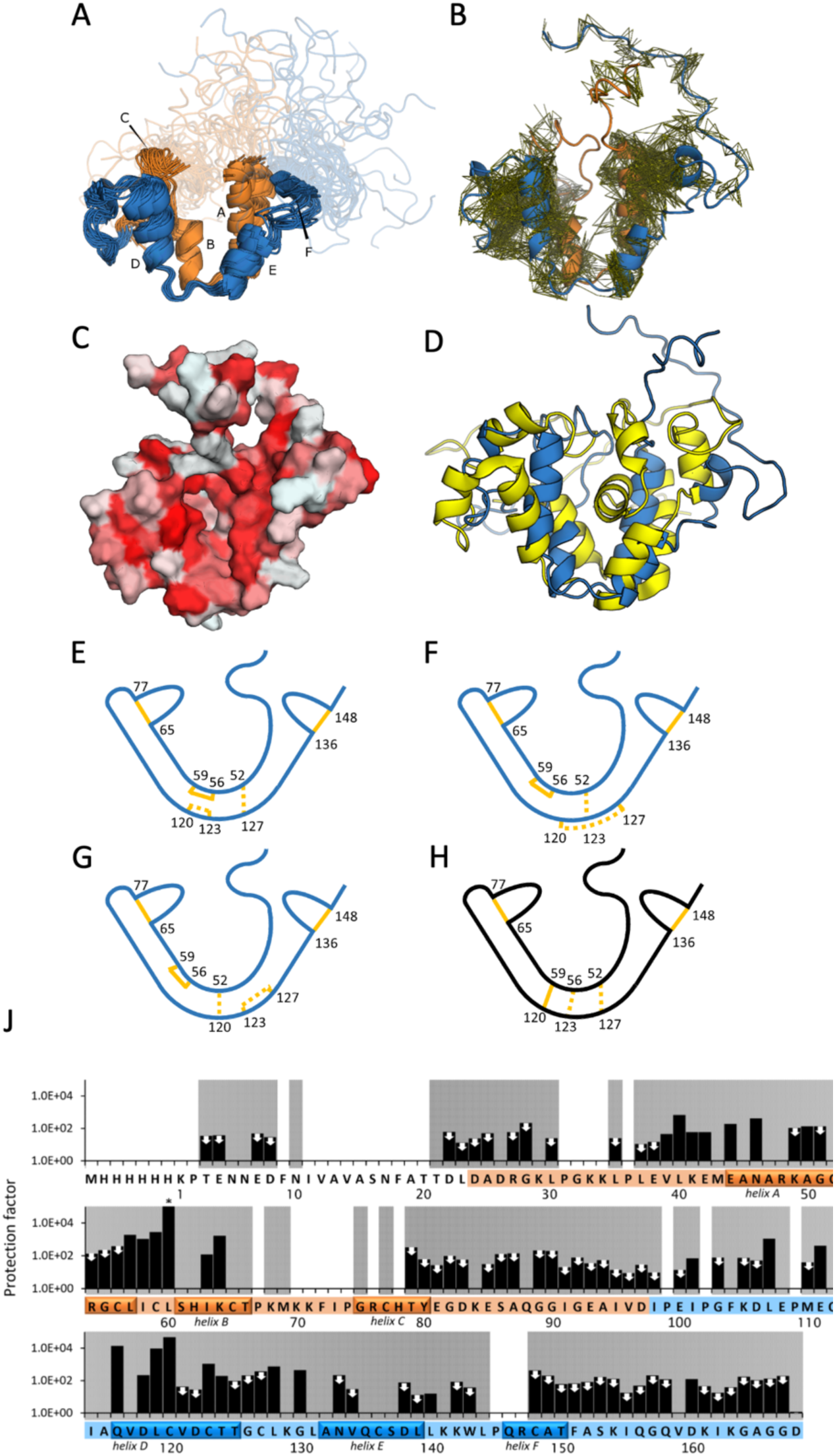
Structural properties of GLuc. **A)** Ensemble of 20 representative structures of GLuc; PDB 8OQB. Residues 1-97 are shown in orange, residues 98-168 are shown in blue, respectively corresponding repeated sequence motifs. Disordered parts are depicted semi-transparently. Small capital letters indicate 6 α-helixes. **B**) Visualization of distance restraints used for the calculation of GLuc’s structure. Each line represents a restraint between two atoms derived from an observed peak in a 3D NOESY spectrum. **C**) Surface representation, where coloring indicates hydrophobicity, from least hydrophobic (white) to most hydrophobic (red) (Eisenberg et al., 1984). **D**) Comparison of 8OQB (blue) and 7D2O (yellow; Wu et al., 2020) **E-G**) Schematic representations of disulfide bond pattern between closely spaced cysteine residues. Bonds confirmed with high confidence by mass spectrometry are shown in solid lines. Three unresolved disulfide patterns are shown as dashed lines. Panel **E** represents the configuration we deem most likely based on MS/MS and structural restraints. **H**) The disulfide bond configuration in 7D2O (Wu et al. (2020)) is not compatible with our data (see SI Figure S11). **J**) Hydrogen-deuterium-exchange rates, determined by NMR. Protection factors were calculated as the ratio of the reference exchange rate in random coil to the measured exchange rate in GLuc’s structure. Two mutually similar regions in the sequence are colored in orange and blue, respectively. White and grey background indicates absence or presence of backbone assignments, respectively. The helices are indicated on the sequence. White arrows mark 72 amides that had exchanged in the dead time, so only a maximum value of the protection factor could be estimated. The asterisk (*) indicates no change detected at the end of the experiment, meaning that the protection factor is likely higher than shown. See also supplementary Figure S12.

### Structural analysis of GLuc by NMR

We realized that the GLuc used for previous structural work included two mutations, E100A and G103R, to increase the expression. Comparing this variant with the wild-type, in our hands, showed less than half the activity (SI Figure S3 and S4). This made us concerned that the mutations might affect the structure as well and led us to undertake an NMR-based structure study with wild-type GLuc.

The ^15^N-HSQC spectrum of GLuc shows a relatively small dispersion of the peaks, indicating that GLuc is less structured than most proteins, although some structured regions are present. We were able to assign 83.5 % of backbone atoms, 72.6 % of side chain hydrogens and 57.7 % of side-chain heavy atoms (supplementary Figure S7 and Figure ST4). Due to broad lines, we were not able to assign parts of the backbone, namely residues 9-20 of the flexible N-terminus, and regions 31-36, 70-78 and 145-147. All missing amide assignments are greyed out in Figure 2J. To calculate the structure we used NOEs, dihedral angle restraints, hydrogen bonds and five disulfide bonds (Figure 2 and supplementary Table S4).

Performing mass spectrometry, we unambiguously assigned three of the five disulfides in GLuc (SI Figure S10). These are C65-C77, C136-C148 and notably including C56-C59 not present in 7D2O. MS analysis was confirmed in two independent labs. We therefore performed individual structure calculations using the software Cyana (Güntert, 2013) with the three possible disulfide configurations for the remaining four Cys residues, i.e.: C52-C127/C120-C123, C52-C123/C120-C127 and C52-C120/C123-127 (Figure 2E-2G). For the three sets of disulfide bonds the target function that Cyana minimizes was 7.0 ± 0.6, 7.3 ± 0.4 and 20.0 ± 0.8, respectively. The structures calculated with the first set of disulfides were able to accommodate the torsion angles calculated from the chemical shifts the best, resulting in fewer violations, and had fewer Van der Waals clashes. Furthermore, we found no evidence of disulfide bond heterogeneity in our NMR spectra and we did not observe multiple peaks for any residues. We therefore choose to use the C52-C127/C120-C123 disulfides for the subsequent structure calculations (Figure 2E).

The C56-C59 disulfide is highly surprising from an evolutionary point of view. The sequence of GLuc likely arose from a gene duplication event (Inouye et Sahara, 2008). Residues 27 to 97 are 27 % identical to residues 98 to 168 and 48 % similar (using the EBLOSUM62 matrix). GLuc’s cysteine spacing is conserved for four out of five cysteine residues in each domain. This suggests that they predate the duplication event, and we expected that the disulfide bonds in which these cysteines participate would be conserved as well. A disulfide bond pattern in line with this hypothesis would connect C52, C56, C65 and C77 (in the first domain) and C123, C127, C136 and C148 (in the second domain). The last two (non-conserved) cysteines would then form a bond with each other (C59-C120) and thus connect these two domains, as suggested previously (Wu et al., 2015). The observation that C59 does not connect to C120 disrupts this pattern.

Our structure model of GLuc is topologically similar to the structure reported by Wu et al., but also features differences. As in the published structure of GLuc-E100A/G103R, our model of native GLuc contains four main α-helices (A, B, D and E), which are arranged pairwise in a double V-shape, where each of the V’s bear some structural similarity, reminiscent of the above mentioned duplication event (Figure 2A). In our structure, the C52-C127 disulfide bond holds the loops at the bottom of the two V’s together (Figure 2F). The disulfide bond that is conserved between the two halves of GLuc’s sequence (C65-C77 in the first half and C136-C148 in the second) stabilizes two small helices (F and C in Figure 2A). These regions are located on relatively unstructured parts of the protein on the open end of the V-shape. In the 7D2O structure, all three disulfides at the bottom of the V’s connect the two halves of the protein (Figure 2J).

We found the N-terminal region (residue 1-27) to be completely disordered, with only a few local NOEs and no long-distance contacts with other parts of the protein. This constitutes a significant difference from the 7D2O where the N-terminus forms a helix between the arms of the V. Considering that this region is not essential for activity (Figure 1D, SI Figure S5), it seems unlikely that it would play an essential role in the assembly of the substrate binding pocket as suggested by Takatsu et al. (2023).

We found a few NOEs between the two arms of the V connecting residue K70 and W143. However, it was not possible to assign these peaks unambiguously because of significant overlap with other close residues with similar shifts. Since the connection was not supported by further NOEs and including it in the structure calculation caused unresolvable violations in other parts of the structure, we chose to leave it out of the final structure calculation. It does however suggest a possible transient interaction between the arms of the V-like structure.

### GLuc is highly dynamic

Even if our structure of GLuc does not reveal a clear substrate binding pocket, we noticed that many residues important for its function are located on helices C and F at the open end of the V, in otherwise unstructured regions. These residues include the catalytic essential residues R76 and R147, which are in conserved positions within the putatively duplicated sections. However, they point away from the suggested substrate groove and the average Cα distance between them is 25 Å (Figure 1E), which requires a major structural rearrangement if they should simultaneously interact with CTZ. The distance was similarly large in 7D2O (21 Å, Figure 1E).

Helices C and F also harbor the aromatic residues F72 and Y80, which interact with CTZ (Larionova et al., 2017). Indeed, two tyrosine residues of *Metridia longa* luciferase, which are equivalent to residues to F72 and Y80 in GLuc, displayed quenching of their intrinsic fluorescence upon substrate binding (Larionova et al., 2017). W143 and L144 residues both cause a luminescence red shift of 3 nm when mutated to valine and alanine, respectively. These shifts were observed for the GLuc variant MONSTA (F72W/I73L/H78E/Y80W). A double mutant shifted by as much as 7 nm compared to the MONSTA background, which brings the total shift of that variant compared to wild-type GLuc to 15 nm (Wu et al., 2016). When histidine 78 is substituted by a glutamate, the light output is three times higher and the wavelength shifts by 4 nm (Kim et al., 2011).

It has been shown that GLuc displays substrate cooperativity (Dijkema et al., 2021; Larionova et al., 2018; Tzertzinis et al., 2012), meaning that its initial rate, as measured by the light intensity, increases with substrate concentration to the power of 1.6-2.0, the Hill coefficient (Hill, 1910). Typically, cooperativity occurs in oligomers with multiple allosteric substrate binding sites (the classic example being hemoglobin (Monod et al., 1965; Stefan & Novère, 2013)).

In this context, it is worth mentioning that a previous study suggested that the two half-domains possess significant activity individually (Inouye & Sahara, 2008). Furthermore, the NOEs in GLuc reveal two almost completely separate hydrophobic regions (Figure 2B) between the arms of the V, which could suggest the existence of two active sites. However, it is difficult to reconcile this with the observation that both R residues can individually knock out the activity completely. Indeed, in our hands, the separate domains have not given rise to luminescence. This, in our view, excludes the presence of two independent binding sites in GLuc.

To assess the local stability of GLuc we performed hydrogen deuterium exchange and calculated protection factors as the ratio between the observed rate and that of the same amino acid in an unstructured peptide. Typically, such factors lie between 10^2^-10^6^ for well-folded proteins (Hughson et al., 1990; Pan & Briggs, 1992; Radford et al., 1992) and in e.g. RNAse A (about ¾ of the length of GLuc) 42 amide protons have protection factors above 10^4^, including 11 above 10^6^ (Wang et al., 1995). As seen in Figure 2J, for GLuc protection factors for only two residues are above 10^4^ and none above 10^6^. In fact, at least 72 of the backbone amides have exchanged before recording the first spectrum, which corresponds to protection factors below approximately 400. GLuc is thus highly dynamic and adapts solvent-exposed conformations much more frequently than typical globular proteins.

The essential residues (R76, R147) and residues that likely interact (K70 and W143) are located far apart from each other in the structure, in regions with low HDX protection factors suggesting that they are highly dynamic. Thus, we propose that these regions cluster around the substrate when it is bound with the bottom of the V forming a hinge (Figure 3). Such motion is also consistent with *kinetic cooperativity* which has been demonstrated most clearly in glucokinase (Porter & Miller, 2012), where the enzyme only adopts its active conformation when the substrate binds. The dynamics between the open and the closed conformation are slow relative to the catalytic turnover. This means that at high substrate concentrations, a new substrate molecule can bind before the active conformation relaxes, thus increasing the efficiency relative to lower substrate concentrations (Qian, 2008). Kinetic cooperativity for GLuc has been suggested before (Larionova et al., 2018) and our results provide a mechanistic explanation supporting this hypothesis.

**Figure 3.**
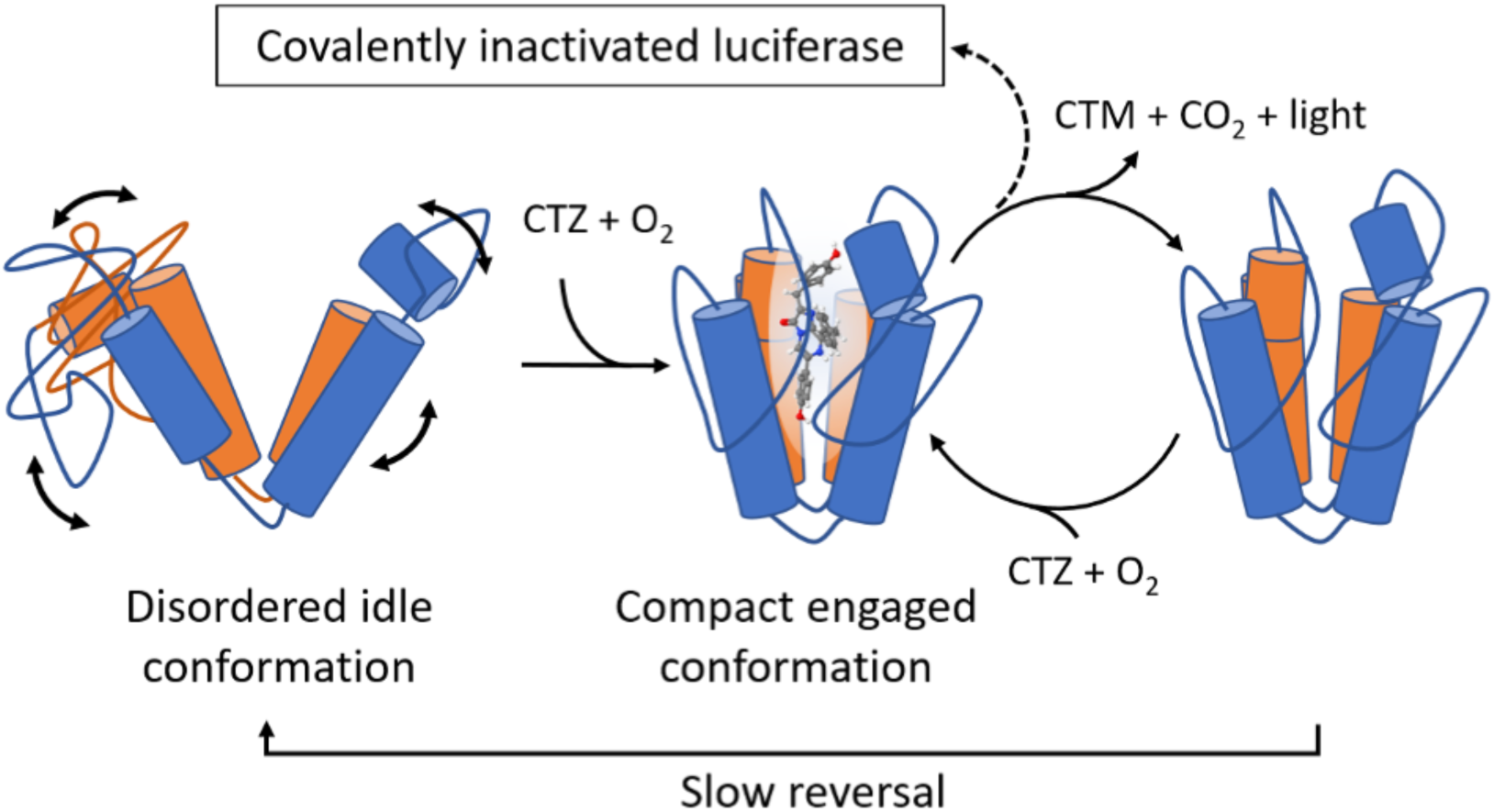
Hypothetical structural changes consistent with kinetic cooperativity. The solution structure represents an open conformation where active site residues are not properly aligned for catalysis (left). On binding of substrate, a major rearrangement takes place which establishes the active conformation (center). After reaction (right), positive cooperativity is observed if, at high substrate concentrations, binding of fresh substrate is more rapid than reversal to the idle disordered state. The catalytic cycle sometimes results in covalent inactivation of the enzyme (top).

In an attempt to capture a closed state, we titrated GLuc with a solution of already oxidized CTZ. In the resulting HSQC spectra, many of the signals disappeared at higher substrate concentrations (SI Figure S13), likely due to conformational exchange at the intermediate timescale. This suggest that the protein may spend a larger fraction of time in the closed state but now with a slower dynamics, supporting the kinetic cooperativity model. Unfortunately, oxidized CTZ did not bind strongly enough fully saturate the substrate-bound conformation and acquire high-quality NMR spectra for a putative closed conformation.

We also hypothesized that the inactivated GLuc obtained after reaction with CTZ might be in a closed form. However, the HSQC spectrum of such inactivated GLuc, was highly heterogeneous and therefore impossible to assign.

The hydrophobic dye 8-Anilinonaphthalene-1-sulfonic acid (ANS) has been used to determine the solvent-accessibility of hydrophobic patches in proteins. We observed strong ANS fluorescence signal at 460 nm for GLuc upon binding, while no signal was observed for a similar concentration of lysozyme, a protein of similar size (SI Figure S8). This can be explained either by the presence of a hydrophobic cavity or by hydrophobic patches in a protein that lacks a well-packed core (Morozova et al., 1995; Qadeer et al., 2012; Sen et al., 2008). Although both models are possible, it does show that GLuc exposes significant hydrophobic surfaces similar to molten-globule states mostly seen under semi-denaturing conditions (Morozova et al., 1995; Sen et al., 2008). Nevertheless, GLuc remains soluble and is stable towards precipitation for weeks at 5°C.

### Substrate reaction is not necessarily accompanied by light production

Furimazine (FRZ, Figure 4g) is an analogue of CTZ (Figure 4F) that is known to produce light with the CTZ-dependent luciferase NanoLuc, but not with GLuc or other members of the copepod luciferase family (Hall et al., 2012; Heise et al., 2013; Li et al., 2021). Because of this, we surmised that FRZ could be a GLuc inhibitor and thus a tool to localize the substrate binding site and lock the closed conformation. We first confirmed that GLuc does not produce light with FRZ and that FRZ did inhibit subsequent reaction with CTZ. However, the onset of this inhibition was not instantaneous which made us suspect that a reaction with the luciferase had taken place. Thus, we performed an experiment in which GLuc was incubated with FRZ in a 20-fold molar excess for up to 24 hours.

**Figure 4.**
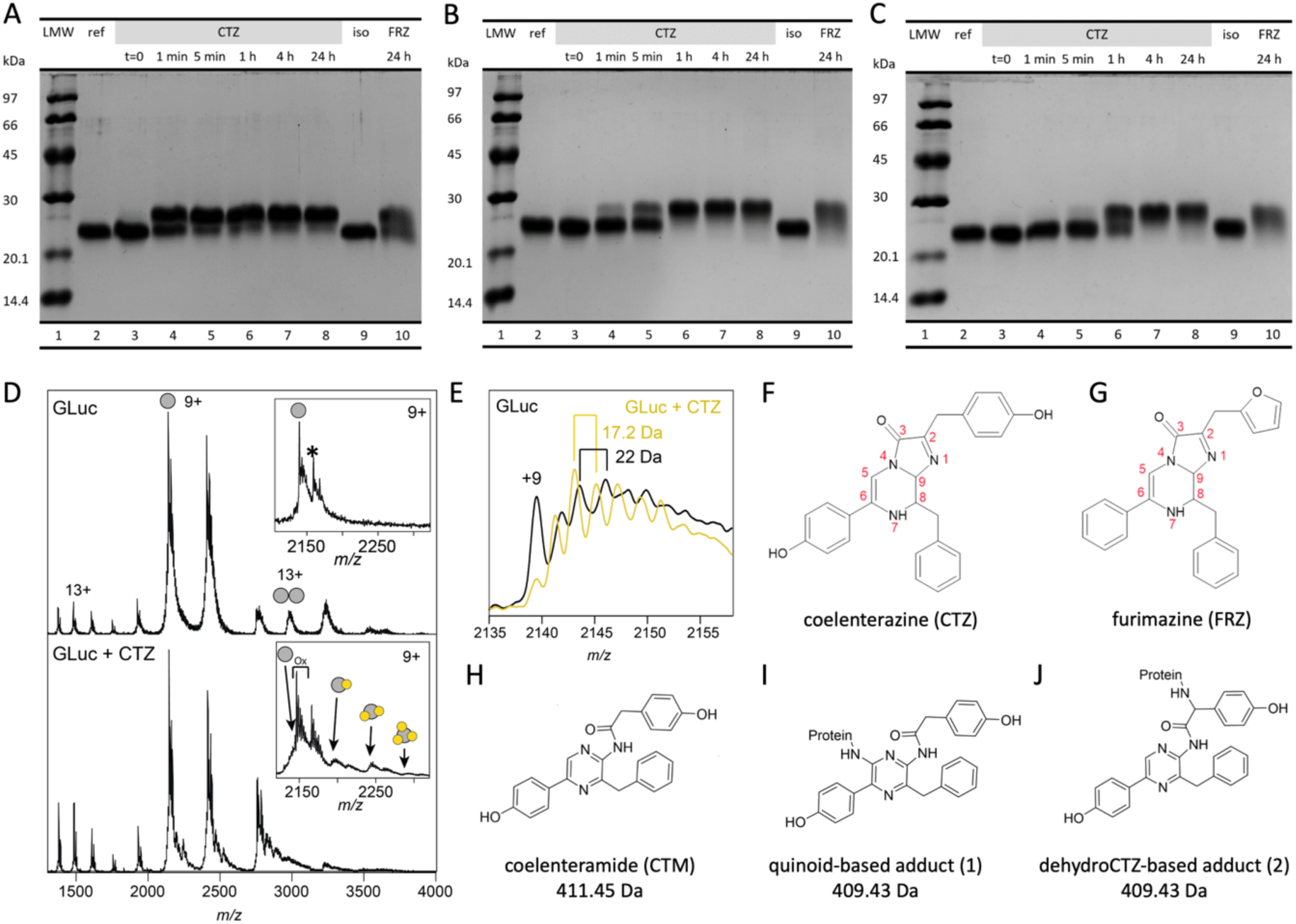
Reaction of wild type, R76A and R147A, respectively, with CTZ and FRZ. **A**) SDS-PAGE of wild-type GLuc before (lane 2) and at indicated time-points after incubation with either coelenterazine (CTZ) (lane 3-8) or furimazine (FRZ) (lane 10) showing formation of adducts. Lane 9 is loaded with a GLuc control, mock-incubated with isopropanol without CTZ for 24 h and lane 1 is a molecular mass ladder. **B** and **C**) As for A but with GLuc-R76A and GLuc-R147A variants, respectively. **D**) ESI-TOF mass spectrum of wild-type GLuc before (top) and after (bottom) incubation with coelenterazine (CTZ). Insert shows a zoom-in on the major peak with up to three adduct species with a progressive mass increase of 409 Da. The mass increase of 178 Da indicated in the reference spectrum by * is likely due to a posttranslational phosphogluconoylation of the His-tag during expression in *E. coli* seen previously (Geoghegan et al., 1999). **E**) The fine structure of the GLuc peaks in the reference mass spectrum (black spectrum) reveals a repeated increase around 22 Da, consistent with sodium adducts. After incubation with CTZ (yellow spectrum), the peaks are separated by approximately 18 Da, suggesting extensive oxygen adducts (see SI Figure S15). **F-J**) chemical structures of coelenterazine (**F**), furimazine (**G**), coelenteramide (**H**) and two possible adducts (**I** and **J**). These adducts can be formed through conversions shown in supplementary figure S14.

Samples were then subjected to SDS-PAGE as we had previously done to study the inactivation of GLuc after its light-emitting reaction with CTZ (Dijkema et al., 2021). Interestingly FRZ-treated GLuc displayed a reduced mobility on SDS-PAGE, which indicated a shift to a higher molecular weight than the untreated luciferase (Figure 4A, lane 10 and SI Figure S9). Since this shift is similar to the one observed for GLuc inactivated by CTZ (Figure 4A, lane 8), we suggest the occurrence of similar but non-light-producing reactions between GLuc and FRZ which lead to covalent adducts visible in SDS-PAGE.

Thus, FRZ is a suicide inhibitor of GLuc as it does not produce light with GLuc, but still reacts and forms adducts. Since excited states can decay non-radiatively through molecular motion (Güsten & Meisner, 1983) we suspect that, contrary to FRZ, CTZ can bind to GLuc and be kept in a rigid conformation in which a photon can be produced. This means that GLuc must have a highly specific interaction with the hydroxybenzyl group attached to carbon 2 of the imidazopyrazinone core of CTZ (Figure 4F), which is replaced by 2-furanylmethyl in FRZ (Figure 4G) or with the hydroxy group on the aromatic ring attached at position 6, which is missing in FRZ.

For inactive variants R76A and R147A, we assayed adduct formation using SDS-PAGE. Both variants were reacted with CTZ and FRZ and were sampled for SDS-PAGE after 1 minute, 5 minutes, 1 hour, 4 hours and 24 hours of reaction (Figure 4B and 4C, lanes 3-8 and 10). As illustrated by Figure 4, the rate of adduct formation is lower in GLuc R76A and GLuc R147A as compared to wild type. However, although their luminescence is less than 10^−5^ of the wild type, adduct formation is still substantial.

Thus, it appears that the absence of luminescence for both substrates with the mutant variants R76A and R147A is mainly caused by a dramatically lower quantum yield compared to the reaction between wild-type GLuc and CTZ. If the low quantum yield is due to a similar inability to bind substrate in a rigid conformation and prevent non-radiative decay of the excited state, this would support the hypothesis that the arginines play an important role in locking CTZ in a rigid conformation.

### Identification of GLuc adducts

To elucidate the inactivation mechanism of GLuc and in an attempt to determine the active site location, we compared GLuc with CTZ-inactivated GLuc by ESI-TOF-mass spectrometry (Figure 4D and E). The spectra of inactivated GLuc showed an adduct corresponding to a mass increase of 409.5 Da. There are two ways (described in Figure S14 and S15) in which an adduct with this mass could be formed: 1) through a reaction between the luciferase and a reactive quinoid entity (M = 409.43 Da, Figure 4i) resulting from a second oxidation of coelenteramide (M = 411.5 Da, Figure 4H) and 2) through addition of present dehydrocoelenterazine (Inouye, 2022) to the luciferase, which would then undergo oxidative carboxylation (M = 409.43 Da, Figure 4J). It is possible that both reactions occur. The occurrence of CTZ-derived quinoids accounts for the irreversible and very fast inactivation of NanoLuc in the presence of hydroxyfurimazine or h-coelenterazine (Coutant et al., 2020). Notably, the inactivation rate of NanoLuc in that study was greatly reduced when substrates incapable of quinoid formation were used. It is likely that the same reaction takes place in GLuc. However, in GLuc, adducts are also formed with FRZ, which is unable to form quinoids. Those adducts must have formed via a different mechanism.

By MS we observed up to three species with increasing molecular mass in steps of 409 Da consistent with a quinoid derivative of CTZ (Figure 4D). This means that either the enzyme stayed catalytically active after the formation of first adducts in situ or that a substantial amount of GLuc-GLuc cross-reaction took place. In subsequent Tandem MS-MS analysis on tryptic digests, no adducts as large as 409 Da were found – likely because quinoid adducts are prone to hydrolysis (Figure S15).

Recording native mass spectra on the unreacted GLuc we found a repeated mass spacing around 22 Da between the fine peaks, which corresponds to the mass of a sodium ion. For reacted luciferase this was superseded by a difference of 16 Da corresponding to oxygen adducts (Figure 4E), suggesting multiple oxidation events. It should be noted that the sample used for these experiments was not completely inactivated and gave an ensemble of modified species probably with different degrees of inactivation. Attempts to obtain a mass spectrum of fully inactivated GLuc were unsuccessful, possibly due to extensive precipitation along with extreme heterogeneity. As extensively described in the supplementary Figure S14, such oxygen adducts can be due to the occurrence of oxygen radicals resulting from a variety of side reactions taking place during CTZ oxidation. Oxygen radicals have indeed been detected in experiments with the CTZ-using Renilla luciferase (Schenkmayerova et al., 2023).

## Concluding remarks

We would like to highlight three main conclusions. Firstly, GLuc is extremely dynamic with little hydrophobic core. Essentially all backbone amide protons exchange rapidly, even the ones in its four main helices. Binding of ANS shows that it also contains large accessible hydrophobic patches which underpins the main findings of the NMR structure. Furthermore, residues that are known to interact with substrate are placed on widely separated and likely dynamic parts of the protein and must come together to bind substrate. This placement of essential substrate-interacting residues on the structure suggests that the structure we determined by NMR represents an idle conformation, which can enter a more structured substrate-bound conformation through a slow exchange. This would explain GLuc’s cooperative behavior through kinetic cooperativity (Porter & Miller, 2012). Secondly, we have identified two substitutions, R76A and R146A, which have dramatic effects on quantum yield in GLuc, but which apparently can still oxidize substrate and with resulting modification. Related to this, although the light-producing reaction is highly substrate-specific for CTZ, the dark reaction is less so. Since GLuc’s ability to produce light from the oxidation of imidazopyrazinone derivatives is limited to CTZ, the 2-hydroxybenzyl side group or OH-group on the phenol ring in position 6 of CTZ must make essential interactions with the protein. This specificity is not compatible with dynamic active site and underpins the folding-upon-binding hypothesis. Finally, up to three adducts are bound to a single GLuc molecule after reaction with CTZ, accompanied by the formation of numerous oxygen adducts. Adduct formation explains the inactivation of GLuc upon reaction. Together our results on GLuc challenges that enzyme function requires a well-defined structure.

## Materials and methods

The composition of the different buffers and media mentioned can be found in the supplementary information (Figure S2).

### Sequence variants and tags

In our previous work on GLuc (Dijkema et al., 2021), we worked with a version that differed from wild type in two positions: E100A and G103R, to improve the expression yield, as described by Wu et al. (2015). This construct furthermore contained an N-terminal 6 histidine tag (HIS-tag) and C-terminal solubility enhancing tag with the sequence GGGDGGGDGGGD (SEP-tag), both separated from the main sequence by a tobacco edge virus (TEV) cleavage site. We used GLuc of wild type sequence for this study. The alanine scan was performed with protein including a HIS-tag and solubility tag, both separated from the main sequence by a TEV cleavage site. GLuc for NMR only had a HIS-tag and no cleavage site. A full comparison of the performance of the three different constructs and the behavior of the resulting proteins can be found in the supplementary information (Figure S3 and S4 and Table S1).

### Large-scale expression and purification

GLuc was heterologously expressed in *E. coli*. Its sequence was codon-optimized for E. coli and placed in a pET-29b(+) expression vector and custom synthesized by TWIST Bioscience. The small-scale expression and analysis of GLuc variants is described below. For the expression, the CyDisCo system (Gaciarz et al., 2016) was used to aid in formation of the correct disulfide bridges as described (Dijkema et al., 2021). Both the CyDisCo plasmid and our expression plasmid were cloned into E. coli strain BL21 DE3. Cells were grown on liquid AB-LB medium (Lauritsen et al., 2011) or minimal medium containing N15-labelled ammonium chloride and C13-labelled glucose (for NMR) to an OD of 0.7, then induced with 1 mM isopropyl-β-D-thiogalactopyranoside (IPTG) and shaken overnight at 37 °C to produce protein. Protein was purified as described by Dijkema et al. (2021). GLuc for Mass Spectrometry or NMR was further purified by size exclusion chromatography using a Superdex 75 column, for other applications it was simply dissolved in and dialyzed against 50 mM Tris buffer pH 8.0 to remove ammonium sulfate leftover from fractionation. Further details can be found in the supplementary information.

### PSSM scores using POSSUM

The wild-type GLuc sequence without a signal sequence or tags was used as a starting point for the collection of sequences by POSSUM (Wang et al., 2017). Similar sequences were collected from the UniRef50 database (Steinegger & Söding, 2018; Suzek et al., 2015). POSSUM was then used to align the sequences, calculate a position-specific scoring matrix (PSSM) and conservation scores for each residue in the sequence. Residues with a conservation score of 0.7 or higher and the ability to form hydrogen bonds were selected for an alanine scan to test their influence on light production. GLuc variants and wild-type reference were produced with a HIS-tag, SEP-tag and TEV sites (see supplementary Figure S3 and Figure ST1). This sequences were codon-optimized for *E. coli* and placed in a pET-29b(+) expression vector and custom synthesized by TWIST Bioscience.

### Small-scale purification of variant analysis

The purification protocol was adapted slightly for working with a large number of variants at a time. After inital collection of the cells, cells were lysed using BugBuster Protein Extaction Reagent®, with the addition of 5 µg mL^−1^ DNAse and 5 mM MgCl_2_. Solid debris was removed by centrifugation, after which protein was collected using immobilized metal affinity chromatography (IMAC) in sequential incubation steps with nickel-coated beads (one time for binding, three times for washing and two times for elution), each time followed by collection of the beads through centrifugation and removal of the supernatant. The protein was concentrated by precipitation in 60 % ammonium sulfate and pellet resuspended in 50 mM Tris buffer, pH 8.0.

### Luminescence assays

Light emission at 480 nm was recorded on a Perkin Elmer LS55 Luminescence spectrometer equipped with a custom-made syringe holder allowing injection of substrate with a closed lid, thermostated at 25 *^◦^*C. After placing the cuvette containing protein into the machine, but before injection of substrate, the level of background light was recorded for 10 s, to be subtracted from the light output later, in data analysis. Substrate was injected into the cuvette while keeping the lid closed and after starting the recording, so that the initial rise in light emission could be captured. Assays and dilutions of GLuc were performed in 50 mM Tris-HCl, at pH 8.0, with 0.1 g L^−1^ bovine serum albumin to prevent the luciferase at very low concentrations from adhering to surfaces. GLuc was kept on ice until use and CTZ was prepared in ice cold isopropanol and was kept dark and cold in a cooling box at −20 *^◦^*C, in a vile with a membrane so that liquid could be taken out without an influx of oxygen. To minimize GLuc inhibition, isopropanol was at most 1 % of the final reaction mixture. The background was subtracted during data analysis and the start of the light signal was set as 0 s.

### Computation of initial rates, decay rates and light integration - LUMPARSER

A custom Python package was used to fit and analyze the data resulting from light assays (https://github.com/FDijkema/LumParser). Using the interface in this package, the background light was subtracted from all datapoints, and the time was set to zero at the start of the appearance of a signal. The signals were then fitted to a double exponential equation to determine decay rates and extrapolated back to t=0 to determine the initial light intensity with minimal error from mixing. Integration was performed by summation of the data points, corrected for the time step of 0.1 s.

### Analysis of adduct formation by SDS-PAGE

Samples were mixed in 50 mM Tris-HCl pH 8.0, to a GLuc concentration of 14 µM and a CTZ concentration of 0.27 mM. The isopropanol concentration was 10 %, to maximize the CTZ concentration without inhibiting the reaction too much. Aliquots were taken out of the reaction mixture and quenched at indicated time points by diluting 21 µL of each sample into 7 µL of 4x concentrated sample buffer for gel electrophoresis with a final concentration of containing 8 % (w/v) sodium dodecyl sulfate (SDS) and 50 mM dithiothreitol (DTT). 1 µL samples were taken from the reaction mixture for activity assays right after.

### NMR spectroscopy

All NMR spectra were recorded on a Bruker Avance III HD 750 MHz spectrometer at 308 K. For structure determination the sample was 0.83 mM ^13^C/^15^N double-labelled GLuc in 50 mM phosphate, pH 6.0, 5 % D_2_O and 0.125 mM sodium trimethylsilylpropanesulfonate (DSS) and 0.02 % sodium azide. HNCACB, HNCACO, HNCO, HBHA(CO)NH, HN(COCA)CB, HCCH-TOCSY, ^13^C-HSQC and ^15^N-HSQC spectra were used for chemical shift assignment. ^1^H-^13^C NOESY-HSQC and ^1^H-^15^N NOESY-HSQC spectra were used for determining NOE distance restraints.

### NMR data analysis and structure prediction

Spectra were processed using TopSpin (Bruker), qMDD (Orekhov et al., 2004) and NMRPipe (Delaglio et al., 1995) and analyzed in CCPNMR Analysis (Vranken et al., 2005). NOEs were used as the main structural restraints, together with dihedral angle restraints based on chemical shifts, calculated with TALOS-N (Shen & Bax, 2015). After manual chemical shift assignment, NOEs were automatically assigned with CYANA (Güntert, 2013). The resulting NOE list was iteratively refined manually using XPLOR-NIH for the structure calculations (Schwieters et al., 2018). In the final rounds of refinement hydrogen bonds were also included as restraints. The 20 lowest energy structures were chosen from the 100 structures in the final calculation to represent the structural ensemble of GLuc.

### Hydrogen-deuterium exchange NMR

Hydrogen/deuterium exchange NMR was performed on ^15^N-labelled freeze-dried GLuc, dissolved in deuterated phosphate buffer with a pD of 6.0 at the start of the experiment. ^15^N-HSQC spectra were recorded at 20 *^◦^*C every 12.7 minutes for more than 22 hours, starting 5 minutes after mixing the sample. We measured the peak heights in the resulting 267 spectra and fitted them to exponential decay curves. The reference decay rates of the respective residues in random coil (Bai et al., 1993) were divided by the experimental decay rates to calculate the protection factors.

### Binding of 8-anilino-1-naphthalenesulfonic acid (ANS)

Fluorescence spectra of GLuc, ANS, GLuc + ANS and lysozyme + ANS were recorded on a Perkin Elmer LS55 spectrometer. The final concentrations were 10 µM for the proteins and 5 µM for ANS, in 50 mM Tris-HCl at pH 8.0. Emission spectra were recorded from 400-600 nm, with a slit with of 5 nm, at an excitation wavelength of 350 nm, with a slit with of 10 nm. Each spectrum was an average of three scans, with a scan speed of 50 nm/s.

### Mass spectrometry for inactivation

Samples were prepared as described above in expression and purification, then buffer exchanged on a spin filter to 1 M ammonium bicarbonate and diluted to the appropriate concentration in the same buffer. For reacted samples, 1 mM CTZ in isopropanol was added to a quarter of the total volume 5 min before recording spectra. High resolution intact mass spectra were acquired on an Orbitrap Fusion (Thermo Fisher Scientific, MA) equipped with an offline nano-electrospray source. The Orbitrap Fusion was operated in intact protein mode. The capillary voltage was 1.5 kV, the transfer tube temperature was maintained at 40 *^◦^*C and the pressure in the ion-routing multipole was 0.011 Torr. Collisional activation was performed by increasing the HCD energy in the ion-routing multipole at a collision energy of 60 V. High-purity nitrogen was used as collision gas. Spectra were recorded using the Orbitrap mass analyzer at a resolution of 60000 in high mass mode and a scan time of 1 ms. Data were analyzed using Xcalibur 3.0 (Thermo Scientific, Waltham, MA).

Native mass spectra were recorded on a Waters Synapt G1 TWIMS MS modified for intact mass analysis (MS Vision, the Netherlands). The instrument was equipped with an offline nanospray source and the capillary voltage was 1.5 kV. The cone voltage was set to 10 V and the source temperature was maintained at 30 *^◦^*C. The source pressure was adjusted to 8 mbar. The ion trap voltage was 10 V and the transfer voltage was 10 V. IM settings were: Wave height 11 V, wave velocity 300 m s^−1^. IMS gas was nitrogen with a flow of 30 mL h^−1^. Data were analyzed using the MassLynx 4.2 Software package (Waters, UK).

### Tandem mass spectrometry

GLuc and CTZ were mixed and left to react for either 10 min or 3 days. Data from the 3 day sample is presented here. These samples, plus a GLuc reference samples were prepared in triplicate. Details on sample preparation can be found in Supplementary Information.

The raw MS files were first searched in Byonic (v3.6.0 from Protein Metrics, Cupertino, CA, USA) using fully specific tryptic cleavage sites (KR) and allowing for 2 missed cleavages against a database consisting only of the GLuc protein sequence including the His tag. Precursor mass tolerance was 10 ppm, and fragment mass tolerance was 20 ppm. No mass recalibration was used. Oxidation at C, M, P, H and W was considered as a variable modification. Additionally, wildcard search was employed (minimum mass of 0 and maximum of 200) considering any residue, which permits identification of unexpected masses at unexpected amino acid residues. Finally, Disulfide option was used also. Maximum precursor mass was 10000, precursor and charge assignments were computed from MS1, maximum number of precursors per MS2 was 2, and smoothing width was 0.01 m/z.

The modifications identified with Byonic were then used as expected modifications in multiple searches by Proteome Discoverer 1.4.1.14 (Thermo Scientific) conducted on the same raw files and same database. Enzyme was trypsin (full). Precursor mass tolerance was 10 ppm and fragment mass tolerance 0.02 Da. Search engine used was Sequest HT with default ion weights. The precursor ions area detector module in Proteome Discoverer was thus used to assign areas, or quantities to the peptides in each sample.

## Acknowledgements

The authors thank Charlotte O’Shea for excellent technical assistance and Yves Janin for numerous helpful discussions and for critical reading of the manuscript. This work was supported by the Novo Nordisk Foundation grant no. NNF19OC0058579 to JRW and the cOpenNMR facility (Novo Nordisk Foundation, NNF18OC0032996) at the Department of Biology, University of Copenhagen.

