## Supplementary material for "A suicidal and extensively disordered luciferase with a bright luminescence"

### Supplementary materials and methods

#### Methods

##### Protein purification

After expression, cells were collected by centrifugation at 8000 g for 30 min. Cells from 1 L culture was resuspended in 20 mL 50 mM Tris, 150 mM NaCl, 10 mM imidazole, pH 8.0. The suspension was sonicated 8 x 30 s at 50 % amplitude while on ice, using a UP200S ultrasonic processor (Hielscer). Solid debris was removed by centrifugation at 15 000 g and 4 °C for 15 min. To remove DNA from the supernatant, 1 % streptomycin was added. After 30 min incubation on ice, precipitated DNA was removed by centrifugation for 30 min at 15 000 g and 4 °C. The clear lysate was loaded onto a gravity-flow column willed with nickel-coated beads, to bind the His-tagged protein through immobilized metal affinity chromatography (IMAC). The column was washed with 50 mM Tris, 150 mM NaCl, 20 mM imidazole, pH 8.0 after which protein was eluted with 50 mM Tris, 150 mM NaCl, 600 mM imidazole, pH 8.0. Peak elution fractions were identified by activity assays and combined. High-molecular weight disulfide-bonded complexes resulted from incorrect disulfide-bond formation were removed through ammonium sulfate fractionation. Clean protein was found in the supernatant at 50 % ammonium sulfate and the pellet at 75 % ammonium sulfate. Gaussia luciferase (GLuc) for Mass Spectrometry or nuclear magnetic resonance (NMR) was further purified by size exclusion chromatography on a GE Healthcare Superdex75 10/30 column, for other applications it was simply dissolved in and dialyzed against 50 mM Tris-HCl, pH 8.0 to remove ammonium sulfate.

##### Tandem MS

Sample concentrations were the following: 0.12 mM GLuc, 0.5 mM CTZ in 32.5 mM ammonium bicarbonate and 10 % isopropanol. After the incubation period, deionized water was added to a total volume of 100 µL and the samples were frozen in liquid nitrogen and lyophilized. Samples then spent several days in transit at room temperature, after which each dried sample (containing approximately 20 µg of total protein) was dissolved in 100 µl of 50mM ammonium bicarbonate, pH 7.8 (Bioultra from Sigma, prepared freshly) and 1 µg of trypsin was added. Incubation proceeded at 37°C for 3h with mild shaking. Afterwards, the samples were dried in a speedvac and re-dissolved in 20 µl of LCMS solvent A (3% acetonitrile, 0.1% formic acid). For each LC-MS/MS run, 1 µl was injected into a C18 trap desalting column (Acclaim pepmap, C18, 3 µm bead size, 100Å, 75 µm x 20 mm, nanoViper, Thermo).

After 6 min of flow at 5µl/min with the loading pump, the 10-port valve switched to analysis mode in which the NC pump provided a flow of 250nL/min through the trap column. The curved gradient (curve 6 in the Chromeleon software) then proceeded from 3% mobile phase B (90% acetonitrile, 5% DMSO, 5% water, 0.1% formic acid) to 45% B in 50min followed by wash at 99% B and re-equilibration. We used a nano EASY-Spray column (pepmap RSLC, C18, 2µm bead size, 100Å, 75µm x 50cm, Thermo) on the nano electrospray ionization (NSI) EASY-Spray source (Thermo) at 60°C. Online LC-MS/MS was performed in DDA (data dependent acquisition) mode using a hybrid Q-Exactive HF mass spectrometer (Thermo Scientific). FTMS master scans with 60,000 resolution (and mass range 150-1350 m/z) were followed by data-dependent MS/MS (30,000 resolution) on the top 5 ions using higher energy collision dissociation (HCD) at 30% normalized collision energy. Precursors were isolated with a 2 m/z window and an isolation offset of 0.5m/z. Automatic gain control (AGC) targets were 1e6 for MS1 and 1e5 for MS2. Maximum injection times were 100ms for MS1 and MS2. The entire duty cycle lasted ~1s. Dynamic exclusion was used with 30s duration. Precursors with charge states 2-7 were included. An underfill ratio of 1% was used.

### Supplementary figures

#### Sequence and conserved cysteine pattern

MKPTENNEDFNIVAVASNFATTDL

DADRGKLPGKKLPLEVLKEMEANARKAGCTRGCLICLSHIKCTPKMKKFIPGRCHTYEGDKESAQGGIGEAIVD

IPEIPGFKDLEPMEQFIAQVDLCVDCTTGCLKGLANVQCSDLLKKWLPQRCATFASKIQGQVDKIKGAGGD

**Figure S1:** Sequence of GLuc, without any expression tags. Two duplicated regions are shown in orange and blue. The non-conserved N-terminal region is shown in grey where the native signal peptide has been replaced with M. The cysteines are highlighted in yellow. Four out of five cysteines form a repeated pattern in each duplicated region.

#### Alignment and conservation

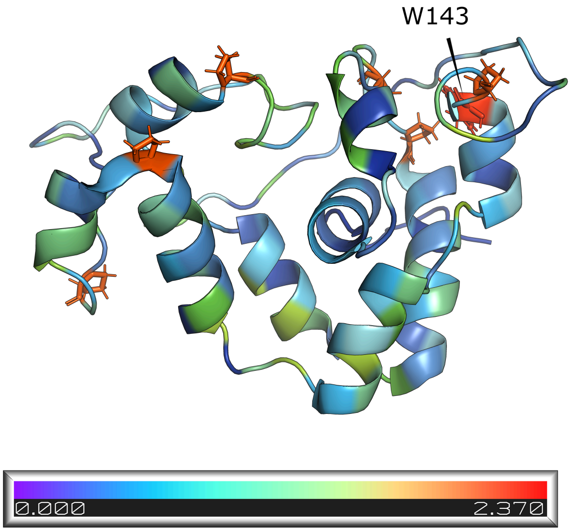


**Figure S2:** The POSSUM conservation score of the residues in GLuc, plotted on its structure.

#### Comparison of activity between differently labelled wild-type variants

Over the course of this project, several different protein sequences were used. In our previous publication (Dijkema et al., 2021), we used a sequence with an N-terminal 6xHIS tag and an C-terminal solubility-enhancing peptide (SEP) tag with the sequence GGGDGGGDGGGD. Both were separated from the main sequence by a TEV cleavage site. The sequence also featured two mutations, E100A and G103R, which have been reported to improve expression (Wu et al., 2015). Later, we tested different constructs to see if they would produce better NMR spectra. We found that both the solubility tag and the mutations decreased the activity.

| | Tag | 1–20 |
|---|---|---|
| SEP-tagged, mutated | M H H H H H H E N L Y F Q G | K P T E N N E D F N I V A V A S N F A T |
| SEP-tagged | M H H H H H H E N L Y F Q G | K P T E N N E D F N I V A V A S N F A T |
| HIS-tagged only | M H H H H H H | K P T E N N E D F N I V A V A S N F A T |

| | 21–60 (30, 40, 50, 60) |
|---|---|
| SEP-tagged, mutated | T D L D A D R G K L P G K K L P L E V L K E M E A N A R K A G C T R G C L I C L |
| SEP-tagged | T D L D A D R G K L P G K K L P L E V L K E M E A N A R K A G C T R G C L I C L |
| HIS-tagged only | T D L D A D R G K L P G K K L P L E V L K E M E A N A R K A G C T R G C L I C L |

| | 61–100 (70, 80, 90, 100) |
|---|---|
| SEP-tagged, mutated | S H I K C T P K M K K F I P G R C H T Y E G D K E S A Q G G I G E A I V D I P **A** |
| SEP-tagged | S H I K C T P K M K K F I P G R C H T Y E G D K E S A Q G G I G E A I V D I P E |
| HIS-tagged only | S H I K C T P K M K K F I P G R C H T Y E G D K E S A Q G G I G E A I V D I P E |

| | 101–140 (110, 120, 130, 140) |
|---|---|
| SEP-tagged, mutated | I P **R** F K D L E P M E Q F I A Q V D L C V D C T T G C L K G L A N V Q C S D L L |
| SEP-tagged | I P G F K D L E P M E Q F I A Q V D L C V D C T T G C L K G L A N V Q C S D L L |
| HIS-tagged only | I P G F K D L E P M E Q F I A Q V D L C V D C T T G C L K G L A N V Q C S D L L |

| | 141–168 (150, 160) | TEV site | SEP tag |
|---|---|---|---|
| SEP-tagged, mutated | K K W L P Q R C A T F A S K I Q G Q V D K I K G A G G D | E N L Y F Q G | D D D G D D D G D D D G |
| SEP-tagged | K K W L P Q R C A T F A S K I Q G Q V D K I K G A G G D | E N L Y F Q G | D D D G D D D G D D D G |
| HIS-tagged only | K K W L P Q R C A T F A S K I Q G Q V D K I K G A G G D | | |

**Figure S3:** Comparison of the sequences of the different GLuc variants used as wild type over the course of this project.

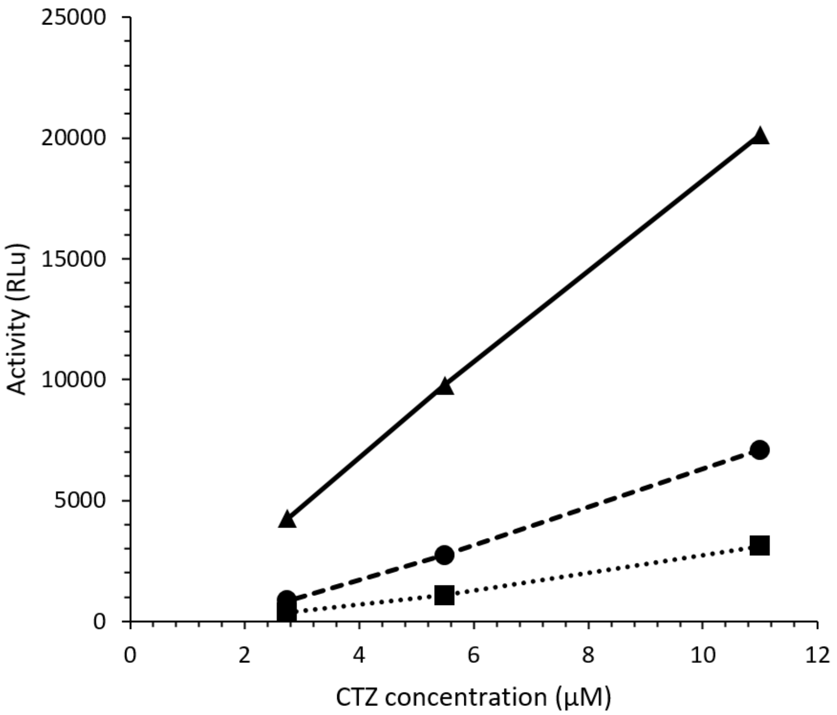


(a)

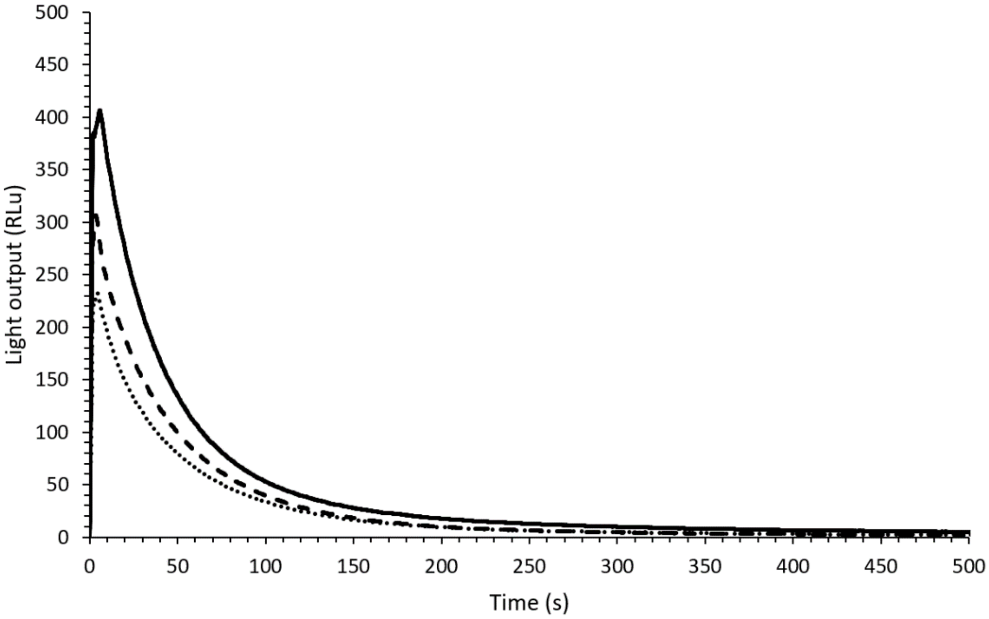


(b)

**Figure S4:** Comparison of the activities of the different GLuc variants used as wild type over the course of this project. Squares/dots represent the tagged and mutated variant, circles/dashes the double tagged variant, triangles/solid line the variant with only a HIS-tag. a) Activity comparison at three different concentrations of CTZ (2.75, 5.5 and 11 µM). The activity was calculated by normalizing the peak height in a light assay by the enzyme concentration. b) Representative light assays at 5.5 µM CTZ (unnormalized). Protein concentrations were 0.23 nM for the tagged and mutated variant, 0.13 nM for the double-tagged variant and 0.041 nM for the variant with only a HIS-tag.

(a)

1 10

Metridia longa luciferase K S T E F D P N I D I V G L E G K F G I

Gaussia luciferase K P T E N N E D F N I V A V - - - - - -

20

T N L E T D L F T I W E T M E V M I K A D I A D T D R A S N F V A

- - - - - - - - - - - - - - - - - - - - - - - - - - - A S N F A T

30 40 50 60

T E T D A N R G K M P G K K L P L A V I M E M E A N A F K A G C T R G C L I C L

T D L D A D R G K L P G K K L P L E V L K E M E A N A R K A G C T R G C L I C L

70 80 90 100

S K I K C T A K M K V Y I P G R C H D Y G G D K K T G Q A G I V G A I V D I P E

S H I K C T P K M K K F I P G R C H T Y E G D K E S A Q G G I G E A I V D I P E

110 120 130 140

I S G F K E M A P M E Q F I A Q V D R C A S C T T G C L K G L A N V K C S E L L

I P G F K D L E P M E Q F I A Q V D L C V D C T T G C L K G L A N V Q C S D L L

150 160

K K W L P D R C A S F A D K I Q K E V H N I K G M A G D R

K K W L P Q R C A T F A S K I Q G Q V D K I K G A G G D -

(b)

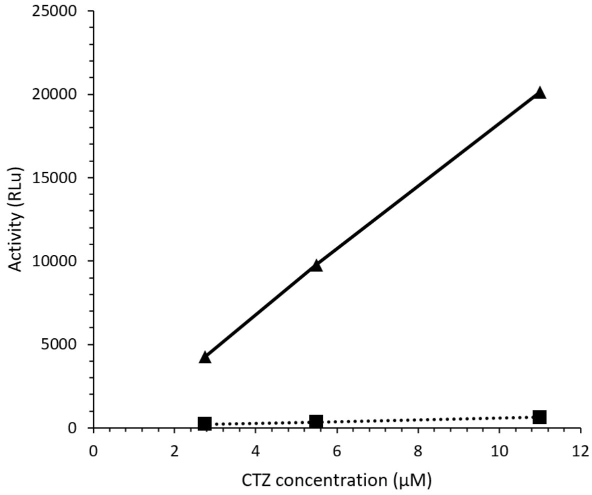


(c)

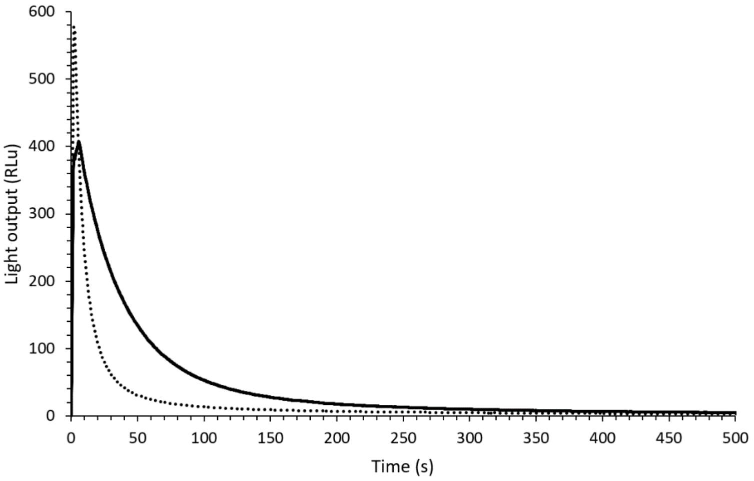


**Figure S5:** Activity of N-terminally truncated GLuc compared to wild type. Squares/dots represent 24-168 N-terminally truncated GLuc, triangles/solid line the full-length variant. Both variants only have a HIS-tag, no SEP-tag or TEV sites (SI Figure S3). a) Sequences of *Metridia longa* luciferase and GLuc with truncation indicated by grey coloring. The N-terminal signal sequences were omitted from both proteins. b) Activity comparison at three different concentrations of CTZ (2.75, 5.5 and 11 µM). The activity at each concentration was calculated by normalizing the peak height in a light assay by the enzyme concentration. c) Representative light assays at 5.5 µM CTZ (not normalized for enzyme concentration). Protein concentrations were 1.17 nM for truncated GLuc and 0.041 nM for full-length GLuc. We note that the inactivation half-life is significantly longer for the 24-168 variant, most likely due to the lower activity.

### Structure determination by NMR

(a)

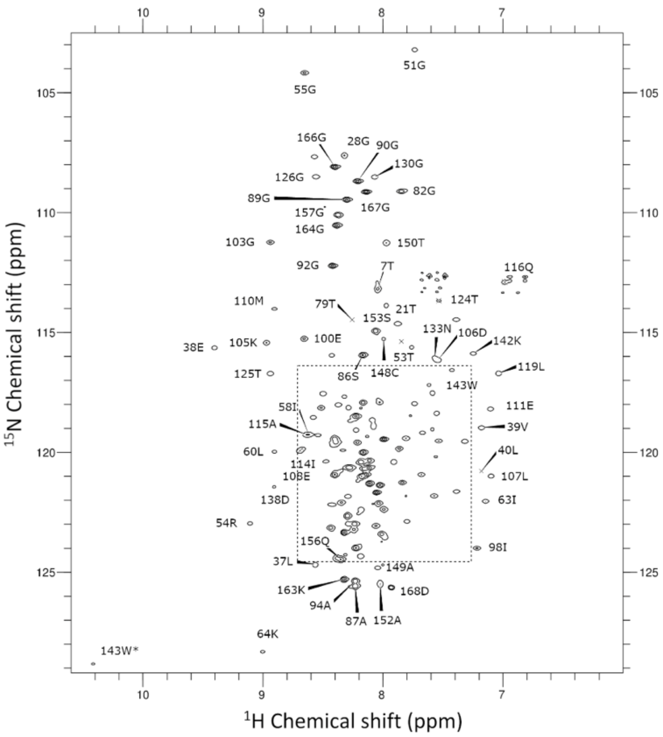


(b)

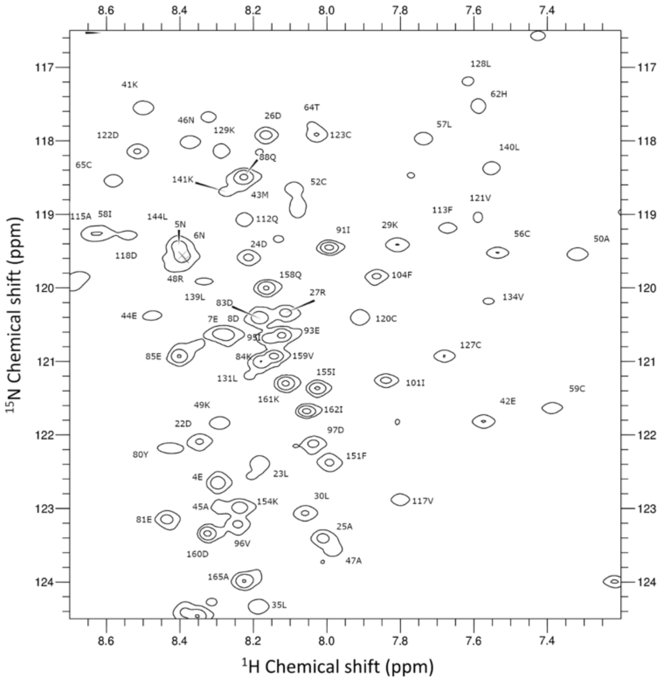


**Figure S6:** a) 2D 1H-15N HSQC of GLuc with backbone assignment. The peaks are labelled with residue numbers and the residue type in single-letter code. See also figure 7.5 for the residue index used. A zoomed-in view of the region marked with a dashed line is shown to better distinguish the peaks (b). In both figures, contours mark an increase of peak intensity of a factor 4. *sidechain resonance

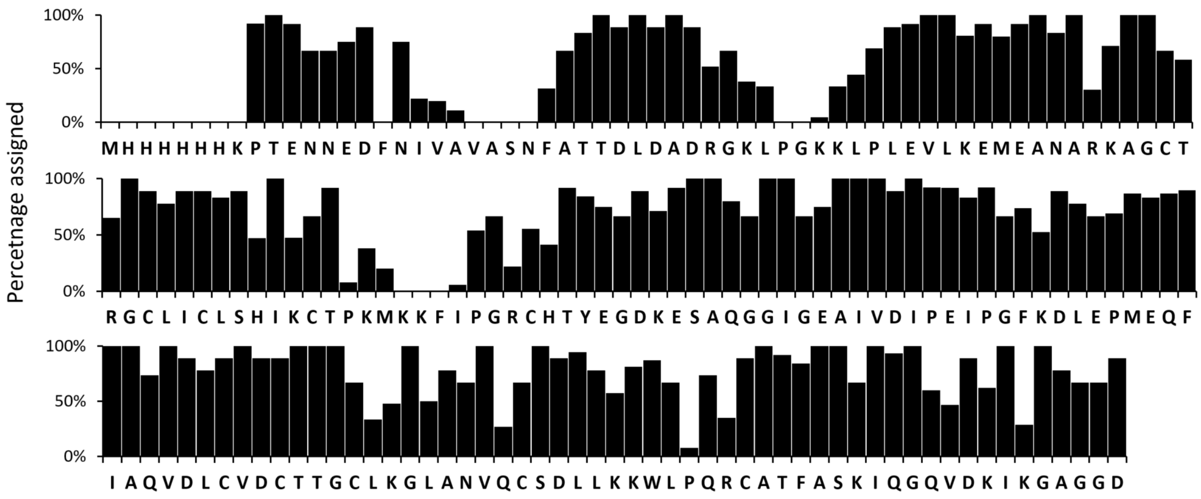


**Figure S7:** Overview of the number of assigned atoms per residue in GLuc.

#### Binding of hydrophobic dye

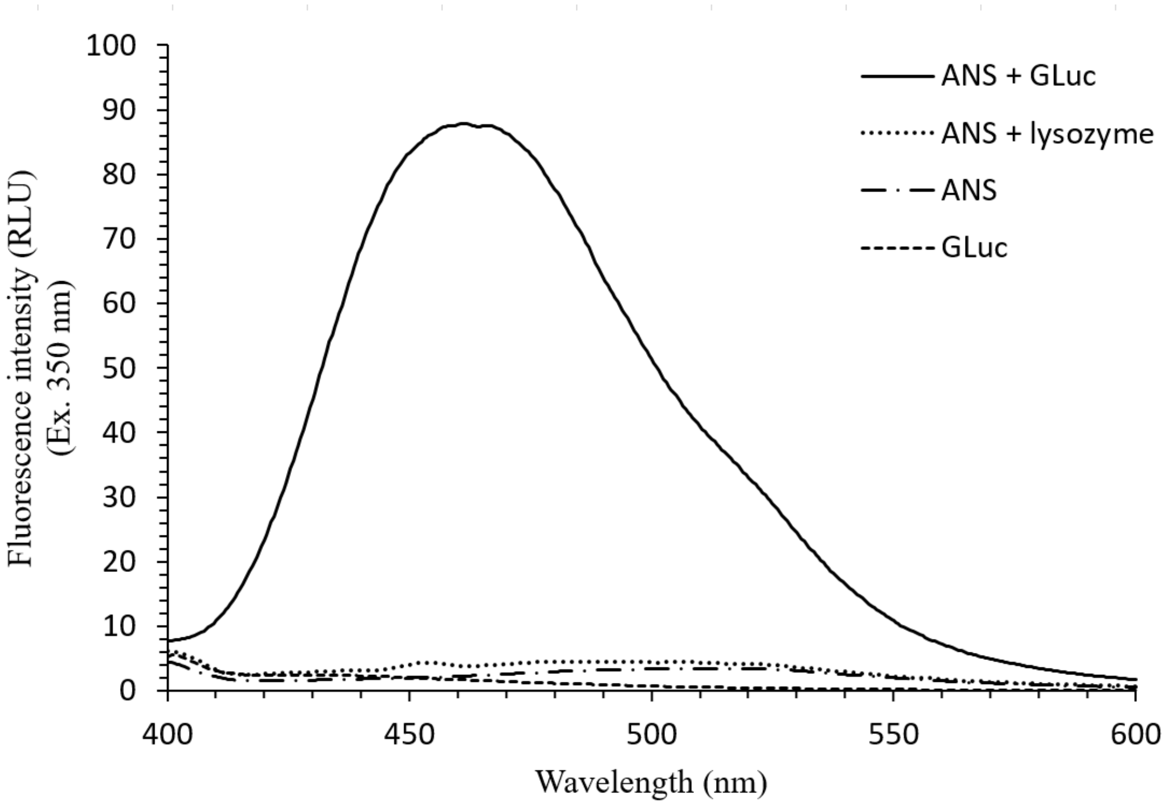


**Figure S8:** Spectrum of 1-Anilino-8-naphthalene sulfonate (ANS) by itself and bound to the internal hydrophobic areas of GLuc.

#### Furimazine modification of GLuc

Although no light is emitted, reaction of GLuc with furimazine results in an upwards shift on a gel, just like with CTZ.

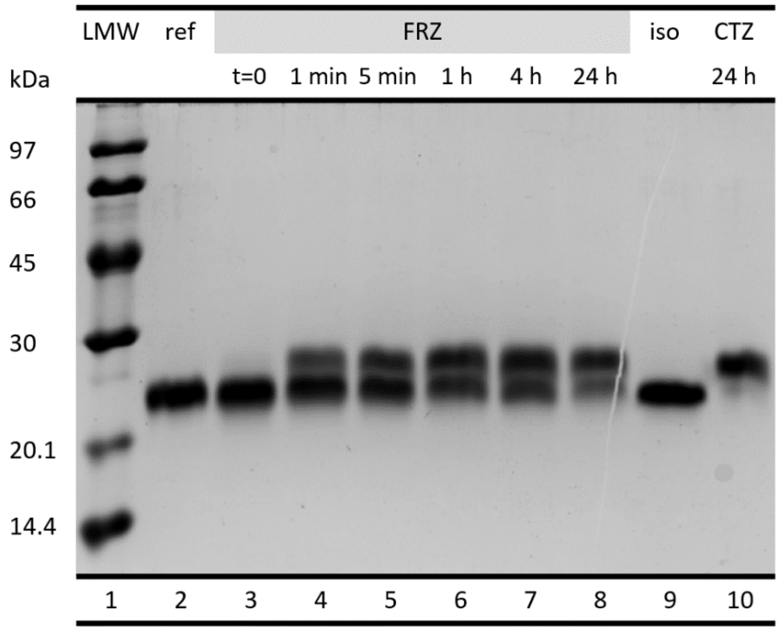


(a) Wild type

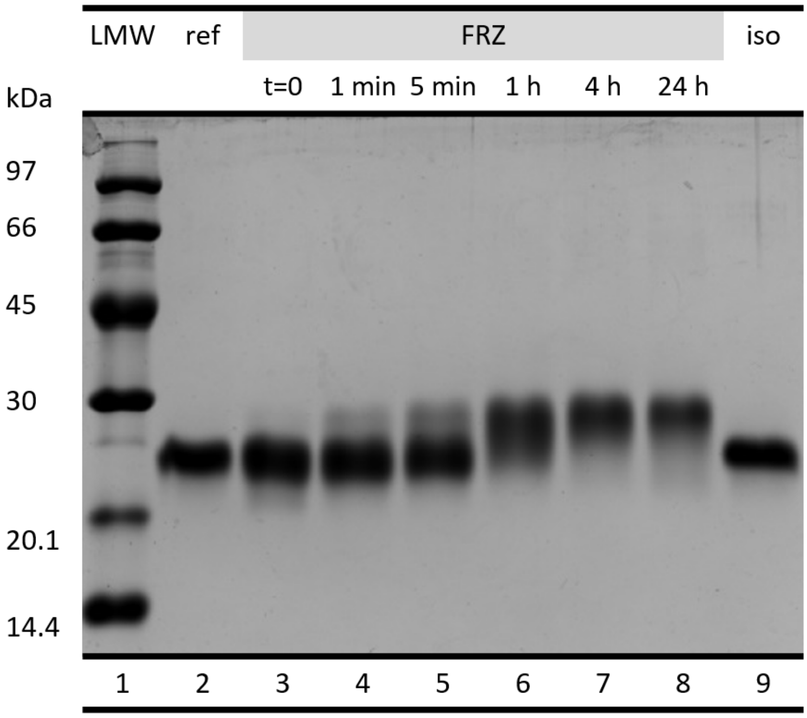


(b) R76A

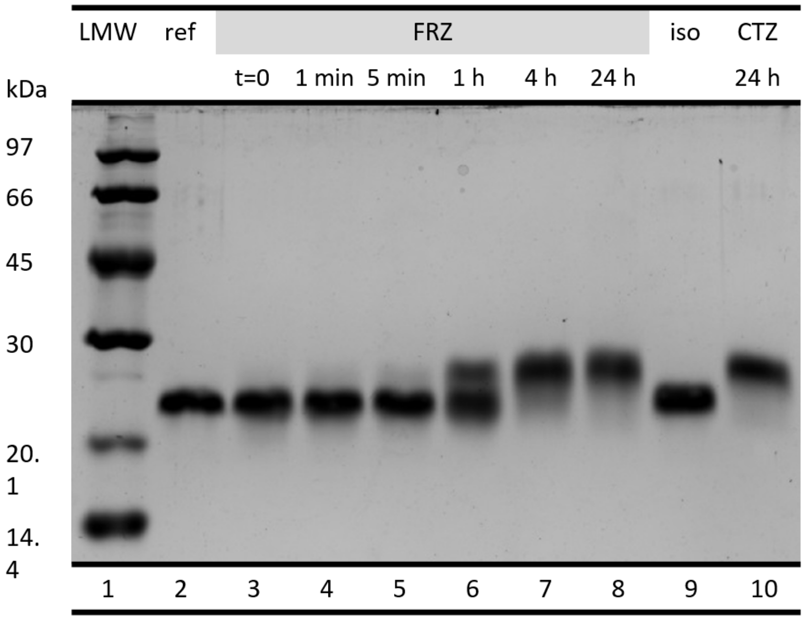


(c) R147A

**Figure S9:** a) After incubation with either furimazine (FRZ) (lane 3-8) or CTZ (lane 10), GLuc shifts upwards on an SDS-PAGE. References of pure GLuc (lane 2) and GLuc incubated with isopropanol (lane 9), the substrate solvent are included. b) Incubation of “inactive” variant GLuc-R76A with frz or CTZ. c) Incubation of “inactive” variant GLuc-R147A with frz or CTZ. Gels were stained with Coomassie brilliant blue.

#### Identification of disulfide bonds

The following peptides were found in in tandem mass-spectrometry on a tryptic digest of GLuc.

| Peptide starts at residue: | Peptide sequence(s) | Residue index of participating cysteines |
|---|---|---|
| 55 | G**c**LI**c**LSHIK | 56, 59 |
| 65<br>77 | **c**TPK<br>**c**HTYEGDKESAQGGIGEAIVDIPEIPGFK | 65, 77 |
| 130<br>148 | GLANVQ**c**SDLLK<br>**c**ATFASK | 136, 148 |
| 106 | DLEPMEQFIAQVDLCVD**c**TTG**c**LK | 123, 127 |
| 106 | DLEPMEQFIAQVDL**c**VD**c**TTGCLK | 120, 123 |

**Figure S10:** Disulfide bonded peptides. Cysteines participating in a disulfide bridge are printed lowercase and bold and highlighted yellow. The last peptide (106) was found with two different disulfide bonds, which means that the pattern is ambiguous.

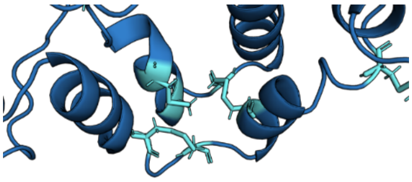

(a)

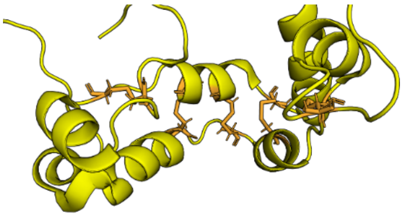

(b)

**Figure S11:** Comparison of the disulfide bond pattern in our calculated structure (top) and 7D2O (Wu et al., 2020). The structured regions in the two respective structures were aligned so that the image is captured from the same angle.

#### Hydrogen-deuterium exchange NMR

To determine which parts of the structure of GLuc were the most rigid, we measured exchange of the backbone amide protons by hydrogen deuterium exchange NMR. Protection factors were determined by dissolving lyophilized protein into deuterated buffer and measuring disappearing of the peaks in an HSQC over time (see methods).

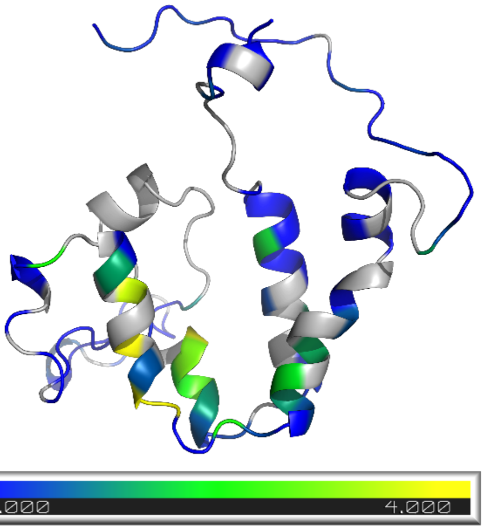


**Figure S12:** Hydrogen-deuterium exchange of backbone amide protons of GLuc. The log of the determined protection factors is plotted on the structure of GLuc here.

#### Changes in the structure upon titration with oxidized CTZ

The binding site of coelenterazine in GLuc is not yet known. We surmised that the reaction product may be similar enough to the substrate that we could use it to bind to GLuc and determine the structure of the bound complex by NMR. Reaction product was generated by leaving CTZ to oxidize in aqueous buffer overnight and the mixture as titrated into an NMR sample of GLuc. HSQC spectra were recorded at increasing concentrations of oxidized CTZ, as seen in figure S11.

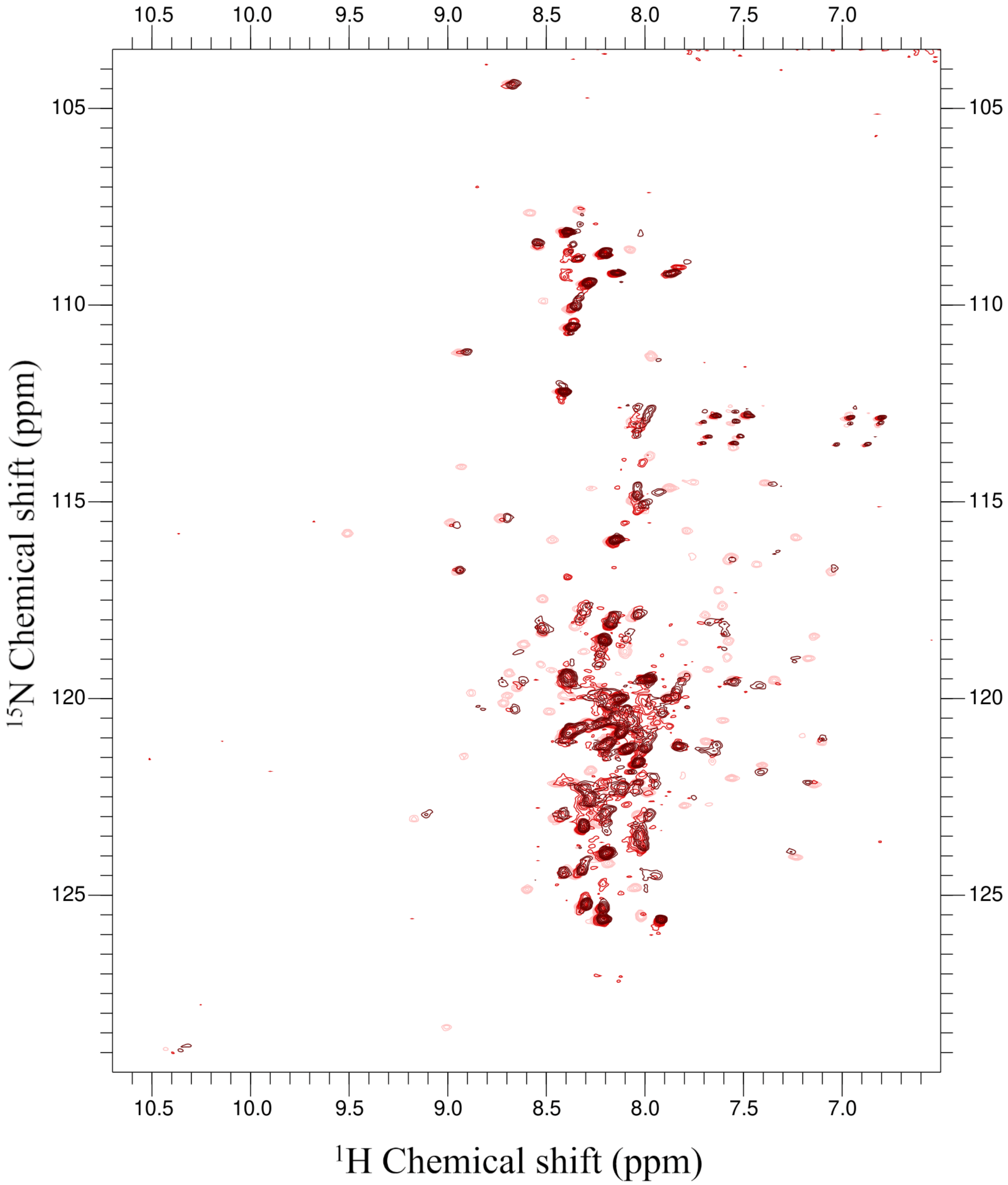


(a) HSQC with oxCTZ titration and change in chemical shifts

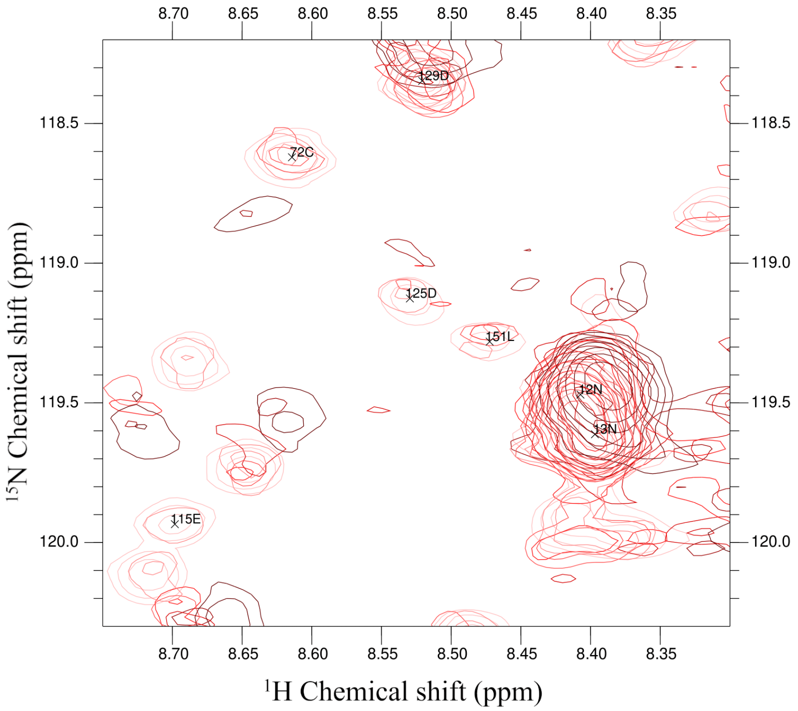

129D
72C
125D
151L
12N
13N
115E
8.70 8.65 8.60 8.55 8.50 8.45 8.40 8.35
118.5
119.0
119.5
120.0
$^{15}$N Chemical shift (ppm)
$^{1}$H Chemical shift (ppm)


(b)

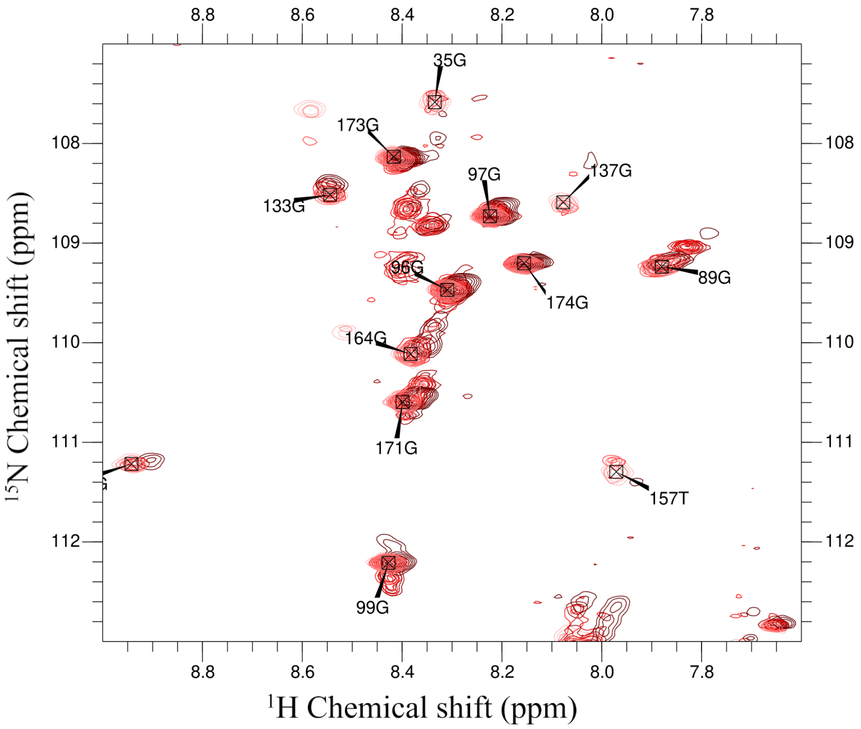

35G
173G
97G
137G
133G
96G
89G
174G
164G
171G
157T
99G
8.8 8.6 8.4 8.2 8.0 7.8
108
109
110
111
112
$^{15}$N Chemical shift (ppm)
$^{1}$H Chemical shift (ppm)


(c)

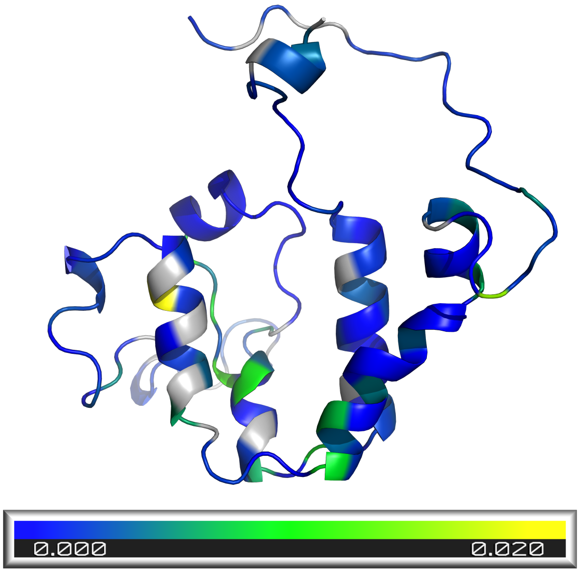


(d)

**Figure S13:** a) HSQC of GLuc, titrated with oxidized CTZ. The oxidized CTZ concentrations were 0, 12, 23, 41, 68, 86 and 190 µM (from light to dark red), which correspond to 0, 0.23, 0.43, 0.78, 1.3, 1.6 and 3.6 equivalents of GLuc protein, respectively. The residue index is shifted by 7 in the HSQC, compared to the index used in the rest of this work, because the HIS-tag was included in the count. Two zoomed areas of the spectrum are shown as examples (b and c) d) Changes in chemical shift at 12 µM CTZ plotted on the native GLuc structure. (d) Change in chemical shifts in ppm (see color bar) after titration with oxCTZ titration visualized on the structure of GLuc in absence of CTZ.

Addition of reaction product to GLuc unfortunately did not lock the structure into a more stable conformation. Changes in chemical shifts were found all over the protein, with most of the larger changes occurring in the helices. Peaks that were not possible to assign or disappeared altogether are indicated in light grey, these are also mostly found in the more structured regions.

Since many peaks could not be assigned or disappeared altogether, it was not possible to determine a structure. Moreover, the binding site could not be determined since changes took place all over the protein.

#### Formation of substrate-derived adducts on GLuc

From coelenterazine (**1**), the main oxidation process starts with a single electron transfer between the anion of coelenterazine and oxygen leading, via the depicted intermediates **2-5**, to the dioxetanone anion (**6**). From this, a decarboxylation will lead to the coelenteramide anion **7** in an excited state which will relax to a ground state either non-radiatively of via the production of a photon. This is then followed by a protonation to give coelenteramide (**12**). The less likely possibilities in this process, such as to a protonation of **6** or **7** before the decarboxylation remain open to discussions as well as the true nature of the photon emitting species. Much of this has been summarized in a recent work on the mechanism of Renilla bioluminescence (Schenkmayerova 2023). Many additional compounds have also been suggested to arise out of additional chemical or enzymatic-based processes. A recent report on the chemistry of the calcium-dependant photoprotein aequorin is providing a quite extensive list of the truly characterized compounds found in the reaction medium and the following is borrowing a lot from this paper (Inouye 2022). Mechanism-wise, a protonation of intermediate **5** to give the hydroperoxide **10** would, upon an elimination reaction also yielding hydrogen peroxide, account for the established occurrence of dehydroCTZ (**16**). From coelenteramide (**12**), a hydrolysis can account for the characterized coelenteramine (**19**) and acid **14**. Moreover, when considering the hydroperoxide **10**, a homolytic break of the would give rise to a hydroxy radical along with radical **17**. A water-based reduction of the latter would lead to alcohol **18** and one more hydroxy radical. The ensuing hydrolysis of **18** would then lead again to coelenteramine (**19**) but also account for the characterized occurrence of the ketoacid **20**. All in all, water peroxide as well as the corresponding hydroxy radical would both explain the occurrence of all the +16 peaks detected by the mass spectroscopy analysis of the CTZ-inactivated GLuc described here.

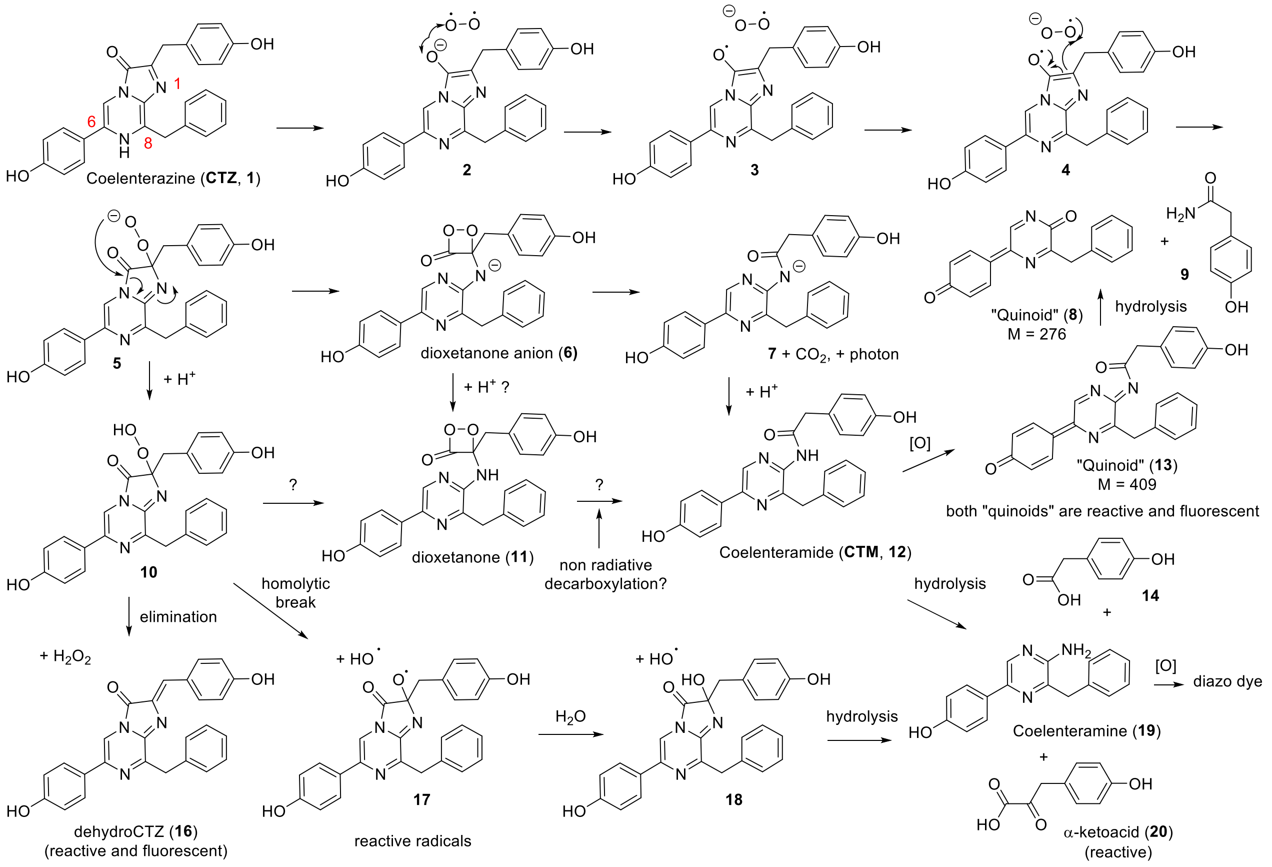


**Figure S14:** various chemicals inferred or characterized occurring upon enzyme-catalysed oxidation of coelenterazine.

Concerning the observed M+409 adducts described in this report, a more hypothetical additional oxidation of coelenteramide (**12**) would give the corresponding and rather reactive quinoid **13** which as depicted in scheme 2, could react with the luciferase. The occurrence of such quinoid and the corresponding enzyme inaction by it is also the most probable explanation for the drastic effects of hydroxyfurimazine or h-coelenterazine on NanoLuc luciferase bioluminescence half-life which changed from hours to minutes (Coutant 2019). Moreover, the hydrolysis of quinoid **13** would lead to the occurrence of quinoid **8** and probably account for the occurrence of the characterized amide **9**. From quinoid **8**, a water-based reduction (not depicted) would then also account for the corresponding (and characterized) hydroxypyrazine.

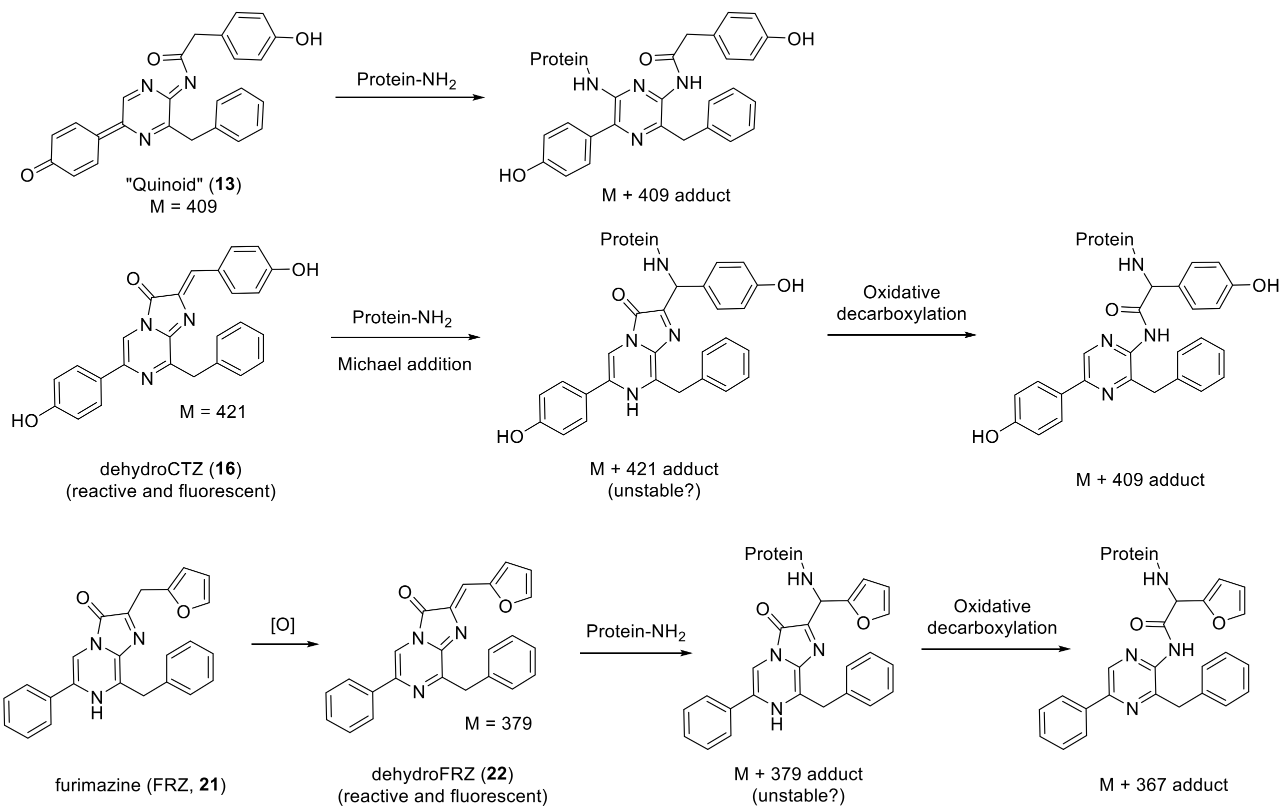


**Figure S15:** possible pathways for the occurrence of protein adducts with coelenterazine or furimazine

As depicted in figure S15, the quinoid **13** is very likely reactive enough to undergo addition reactions with nucleophilic components of the luciferase and thus lead to the observed M+409 adducts. However, another more general mechanism can be suggested. In fact, dehydroCTZ (**16**) can undergo a Michael addition with the same nucleophilic components and this was demonstrated in the past (Tanaka 2009). The resulting covalently bonded imidazopyrazinone is then likely to undergo a chemical (or, less likely, enzymatic) oxidative decarboxylation to give, in the case of CTZ to a protein adduct with the same additional 409 mass. The oxidative decarboxylation of dehydroCTZ (**16**) covalently bonded to the photoprotein symplectin has actually been suggested to explain *Symplectoteuthis* squid bioluminescence (Isobe 1998). Moreover, pholasin luciferin is also made of a protein covalently bonded to dehydroCTZ (**16**) and its resulting bioluminescence is radical triggered (Tanaka 2009). When considering furimazine (FRZ, **21**), a lack of a properly placed hydroxy group makes it rather unlikely that any process would lead to quinoid species. On the other hand, the occurrence of dehydrofurimazine (**22**) is very plausible and then, as depicted in scheme 2 a similar pathway would also lead to protein adducts with an additional mass of 367 Da in this case.

| GLuc type | [CTZ] (µM) | Initial act. (RLU) | Specific act. (RLU/nM) | Ratio | $k_1$ | $k_2$ | Rel. con. $k_1$ |
|---|---|---|---|---|---|---|---|
| SEP-tagged, mutated | 11 | 708 | 3117 | 0.15 | 0.048 | 0.0012 | 0.50 |
| | 5.5 | 244 | 1074 | 0.11 | 0.023 | 0.0027 | 0.75 |
| | 2.75 | 80 | 353 | 0.08 | 0.010 | 0.0026 | 0.62 |
| SEP-tagged | 11 | 914 | 7088 | 0.35 | 0.050 | 0.0015 | 0.54 |
| | 5.5 | 353 | 2735 | 0.28 | 0.023 | 0.0028 | 0.80 |
| | 2.75 | 109 | 848 | 0.20 | 0.010 | 0.0025 | 0.67 |
| HIS-tagged only | 11 | 819 | 20120 | 1.0 | 0.031 | 0.0017 | 0.44 |
| | 5.5 | 398 | 9776 | 1.0 | 0.024 | 0.0028 | 0.67 |
| | 2.75 | 173 | 4255 | 1.0 | 0.016 | 0.0024 | 0.59 |

**Table S1:** Comparison of the activity and decay kinetics of different GLuc sequences. Light assay decay slopes were fitted to double exponential equations to calculate the decay rate constants. The protein concentrations were 0.23 nM for the tagged and mutated variant, 0.13 nM for the double-tagged variant and 0.041 nM for the variant with only a HIS-tag. Initial act. = activity (determined by light peak height in relative light units), Specific act. = specific activity calculated by normalizing activity to protein concentration, ratio = specific activity relative to the activity of GLuc with only a HIS-tag at the same CTZ concentration, k1 = decay rate constant 1, k2 = decay rate constant 2, Rel. con. = relative contribution.

| Variant | Rel. act. | $k_1$ | $k_2$ | Rel. contr. $k_1$ | Rel. Q.Y. |
|---|---|---|---|---|---|
| H78A | 1.71 | 0.046 | 0.004 | 0.79 | 1.07 |
| D24A | 1.26 | 0.022 | 0.004 | 0.75 | 1.05 |
| N133A | 1.08 | 0.035 | 0.003 | 0.77 | 0.93 |
| D122A | 1.03 | 0.029 | 0.003 | 0.76 | 1.28 |
| E42A | 1.02 | 0.014 | 0.004 | 0.59 | 0.47 |
| WT | 1.00 | 0.025 | 0.004 | 0.73 | 1.00 |
| D83A | 0.47 | 0.038 | 0.005 | 0.79 | 0.24 |
| D97A | 0.45 | 0.039 | 0.005 | 0.78 | 0.35 |
| D118A | 0.36 | 0.026 | 0.004 | 0.79 | 1.11 |
| N46A | 0.32 | 0.019 | 0.004 | 0.72 | 0.47 |
| E44A | 0.040 | 0.032 | 0.008 | 0.54 | 0.11 |
| R54A | 0.027 | 0.033 | 0.009 | 0.97 | 0.07 |
| Y80A | 0.012 | 0.049 | 0.014 | 0.63 | 0.03 |
| R76A | 2.3E-5 | - | - | - | - |
| R147A | 1.4E-5 | - | - | - | - |
| D168A* | - | 0.027 | 0.004 | 0.77 | - |
| R27A* | - | 0.027 | 0.004 | 0.68 | - |
| 1-151 | 0.0013 | - | - | - | - |
| 1-155 | 0.014 | - | - | - | - |
| 1-164 | 0.19 | - | - | - | - |
| 24-168 | 0.035 | 0.059 | 0.0031 | 0.61 | - |

**Table S2:** The relative activity of variants of GLuc was measured through measuring the peak height in a light assay and dividing by the wild type activity. The light assay decay slopes were fitted to double exponential equations to calculate the decay rates. Alanine substitution variants were measured in a double-tagged background (Figure S3) and compared to a double-tagged wild-type reference. In these assays the CTZ concentration was 5 µM. Truncation variants were measured in a background with only a HIS-tag (Figure S3) and compared to wild-type reference with only a HIS-tag. In these assays the CTZ concentration was 5.5 µM. *D168A and R27A expressed poorly, which resulted in a purified concentration that was too low to determine. Rel. act. = relative activity, k1 = decay rate constant 1, k2 = decay rate constant 2, Rel. contr. = relative contribution, Q.Y. = quantum yield.

Activity and decay of N-terminal truncated GLuc

It has been reported that removal of the first 72 N-terminal residues of the homolog of GLuc in copepod species *Metridia longa* results in a 10-fold increase in luminescence (Markova et al., 2012). We have investigated if the analogous truncation in GLuc had a similar effect, but as figure S5 and table ST3 show, the activity dropped about 20- fold instead. Additionally, the decay rate increased, meaning that the truncated variant inactivated faster than wild type as well as being less active.

| GLuc type | [CTZ] (μM) | Initial act. (RLU) | Specific act. (RLU/nM) | Ratio | $k_1$ | $k_2$ | Rel. con. $k_1$ |
|---|---|---|---|---|---|---|---|
| FL | 11 | 819 | 20120 | 1.0 | 0.031 | 0.0017 | 0.44 |
| | 5.5 | 398 | 9776 | 1.0 | 0.024 | 0.0028 | 0.67 |
| | 2.75 | 173 | 4255 | 1.0 | 0.016 | 0.0024 | 0.59 |
| 24-168 | 11 | 751 | 642 | 0.031 | 0.066 | 0.0032 | 0.58 |
| | 5.5 | 402 | 344 | 0.035 | 0.059 | 0.0031 | 0.61 |
| | 2.75 | 250 | 214 | 0.050 | 0.045 | 0.0054 | 0.65 |

**Table S3:** Comparison of the activity and decay kinetics N-terminally truncated GLuc (24-168) compared to full length protein (FL). Light assay decay slopes were fitted to double exponential equations to calculate the decay rate constants. Initial act. = activity (determined by light peak height in relative light units), Specific act. = specific activity calculated by normalizing activity to protein concentration, ratio = specific activity relative to the activity of FL GLuc at the same CTZ concentration, k1 = decay rate constant 1, k2 = decay rate constant 2, Rel. con. = relative contribution.

| **Noe distance restraints (upper-bound)** | | |
|---|---|---|
| Short-range | \|i-j\|=0 | 255 |
| Medium-range | 1<=\|i-j\|<=4 | 919 |
| Long-range | \|i-j\|>4 | 290 |
| All | | 1464 |
| *Dihedral angle* | | 172 |
| *Hydrogen bonds* | | 7 |
| *Disulfide bonds* | | 5 |
| **Average pairwise RMSD (Å)** | | |
| Residue 34-81, 95-134 | | |
| Backbone CA | | 5.1 |
| All backbone atoms | | 7.9 |
| Heavy atoms | | 8.7 |
| **Ramachandran plot** | | |
| 20 structures average | | |
| Residues in most favored regions | | 73.90 % |
| Residues in additionally allowed regions | | 7.50 % |
| Residues in generously allowed regions | | 1.10 % |
| Residues in disallowed regions | | 1.00 % |
| Total number of residues | | 127 |
| **Noe violations** | | |
| 20 structures average | | |
| violations between 0.1 and 0.2 | | 44.2 |
| violations between 0.2 and 0.3 | | 8.15 |
| violations greater than 0.3 | | 2.15 |

**Table S4:** Overview of statistics for the structure calculation of GLuc. CYANA was used for peak assignment and initial structure calculations. The final structure calculation was performed using XPLOR-NIH. 100 random seed structures were initiated and optimized, after which the 20 structures with the lowest energy were selected for the final ensemble. Residues 28-156 were included in the calculation. Only residues in structured regions were included for the calculation of the RMSD.
